# Controlling and co-ordinating chitinase secretion in a *Serratia marcescens* population

**DOI:** 10.1101/652685

**Authors:** Marília de Assis Alcoforado Costa, Richard A. Owen, Triin Tammsalu, Grant Buchanan, Tracy Palmer, Frank Sargent

## Abstract

*Serratia marcescens* is a γ-Proteobacterium and an opportunistic animal and insect pathogen. The bacterium exhibits a complex extracellular protein ‘secretome’ comprising numerous enzymes, toxins and effector molecules. One component of the secretome is the ‘chitinolytic machinery’, which is a set of at least four chitinases that allow the use of insoluble extracellular chitin as sole carbon and nitrogen source. Secretion of the chitinases across the outer membrane is governed by the *chiWXYZ* operon encoding a holin/endopeptidase pair. Expression of the *chiWXYZ* operon is co-ordinated with the chitinase genes and is also bimodal, since normally only 1% of the population expresses the chitinolytic machinery. In this work, the role of the ChiR protein in chitinase production has been explored. Using live cell imaging and flow cytometry, ChiR was shown to govern the co-ordinated regulation of *chiWXYZ* with both *chiA* and *chiC*. Moreover, overexpression of *chiR* alone was able to increase the proportion of the cell population expressing chitinase genes to >60%. In addition, quantitative label-free proteomic analysis of cells overexpressing *chiR* established that ChiR regulates the entire chitinolytic machinery. The proteomic experiments also revealed a surprising link between the regulation of the chitinolytic machinery and the production of proteins involved in the metabolism of nitrogen compounds such as nitrate and nitrite. The research demonstrates for the first time that ChiR plays a critical role in controlling bimodal gene expression in *S. marcescens*, and provides new evidence of a clear link between chitin breakdown and nitrogen metabolism.

**IMPORTANCE:** The opportunistic pathogen *Serratia marcescens* secretes chitinases through the action of the *chiWXYZ* operon, which encodes a holin/endopeptidase pair. Expression of *chiWXYZ* is normally bimodal, with only 1% of the population transcribing these genes. In this work, it is demonstrated that overexpression of *chiR* induces exquisitely co-ordinated holin/endopeptidase and chitinase gene expression in the majority of the population of cells, establishing that ChiR is a key player in biomodal gene expression. This work also reveals for the first time that co-operating pathways are induced by ChiR, including enzymes involved in ammonia, nitrite and nitrate metabolism. This work expands knowledge of basic bacterial physiology and could have applications in the biomedical and biotechnological research fields.

## INTRODUCTION

Bacterial cells are exposed to a range of environmental conditions and external signals. To adapt and survive, they must sense their environment, integrate the incoming information and adjust their metabolism accordingly. Protein secretion systems play critical roles in sensing and adaptation pathways, allowing bacteria to compete with other microbes; to interact with and manipulate host cells; or to unlock insoluble or complex nutrient sources. *Serratia marcescens* is a γ-Proteobacterium and some strains are known for production of the red secondary metabolite prodigiosin (1). The bacterium is an opportunistic pathogen, responsible for 1.4% of healthcare acquired infections (2, 3) and strains that are also insect pathogens (for example DB10 and its derivative DB11 (4)) have proven to be rich model systems for understanding the basic physiology of the organism (5).

*S. marcescens* is a prolific secretor of proteins with a diverse ‘secretome’ including haemolysin, phospholipases, proteases, and various toxins and effector molecules. A major component of the secretome is the ‘chitinolytic machinery’, which is a set of enzymes that allow the use of chitin as sole carbon and nitrogen source (6). The four known chitinases of *S. marcescens* are: ChiA, an exochitinase that attacks the polymer from the reducing end; ChiB, an exochitinase that recognises the nonreducing end; ChiC, an endochitinase that performs internal cleavage of the biopolymer; and Cbp21, which is a copper-dependent lytic polysaccharide monooxygenase (LPMO). All of these enzymes are found outside the cell and this location is essential for their physiological function (6).

The *S. marcescens* DB10 strain is an insect pathogen and a tractable model organism for the study of protein secretion (5). A random mutagenic screen identified two genetic loci, *chiWXYZ* and *chiR*, that were important for chitinase secretion (7). The *chiWXYZ* operon is distantly related to a bacteriophage lambda lysis cassette and encodes a holin-like protein (ChiW) and an L-Ala D-Glu endopeptidase termed ChiX (7, 8). Mutant strains deleted for *chiW* or *chiX* were found to be defective in secretion of the chitinolytic machinery with the enzymes accumulating in the periplasmic compartment (7).

Early work in the *S. marcescens* 2170 strain identified ChiR as being important for the production of chitinase activity (9). ChiR is a member of the <u>L</u>ysR-<u>T</u>ype <u>T</u>ranscriptional <u>R</u>egulator (LTTR) family, which is a widespread group of DNA binding proteins (10). The LTTR family has been found to regulate the expression of a variety of genes involved in metabolism, virulence, quorum sensing and motility. Deletion of the gene encoding *chiR* in *S. marcescens* has been reported to result in a drop in extracellular chitinase activity, while supply of extra copies of *chiA* on a plasmid resulted in a relative increase in transcripts from the chitinase genes (11).

Experiments with *S. marcescens* strains carrying chromosomal gene fusions encoding fluorescent reporter proteins demonstrated that expression of the *chiWXYZ* operon was co-ordinated with expression of the *chiA* gene (7). Moreover, these experiments revealed that expression was bimodal with only ~1% of the total cellular population actively producing the chitinase and its secretion system (7). This bimodal expression pattern adds an extra level of complexity that complicates the discovery of new secretion pathway components or secreted substrates.

In this work, the role of ChiR in controlling the co-ordinated bimodal expression of the chitinolytic machinery genes has been explored. Live cell imaging and flow cytometry approaches demonstrated that increased cellular levels of ChiR induced expression of *chiA, chiC* and *chiX* in the majority of the population of cells. Quantitative label-free proteomic analysis was then used to compare the cellular proteome of a Δ*chiR* strain with one overproducing ChiR. This approach revealed that ChiR was not only critical to the production of the chitinolytic machinery, but also had a role in preparing the cells to metabolise the products of chitin breakdown. Thus, ChiR was implicated in the regulation of the assimilatory NADH-dependent nitrite reductase, a nitrite/nitrate antiporter and the energy-conserving respiratory nitrate reductase.

## RESULTS

### A GFP-ChiC fusion demonstrates bimodal expression of the *chiC* gene

ChiC is an endochitinase widespread in Gram-negative bacteria (12, 13). To investigate the native expression pattern of *chiC* in *S. marcescens* DB10, a mutant strain was constructed that would encode a translational fusion between Green Fluorescent Protein (GFP) and ChiC. The construct was placed at the native *chiC* locus on *S. marcescens* chromosome under native transcriptional regulation. The resultant *S. marcescens* strain was named MC03 (as DB10, ϕ*gfp::chiC*, Table 1) and production of the GFP-ChiC fusion protein was then assessed by fluorescence microscopy (Figure 1). The MC03 strain was incubated for 16 hours in either rich or minimal growth media, diluted and added to an agarose microscope mounting slide. The cells were then observed by microscopy using DIC (differential intensity contrast) for cell visualization and a FITC (fluorescein isothiocyanate) filter for GFP detection. Microscopic observations clearly revealed a relatively low number of fluorescent cells (referred to as ‘ON’ with regard to expression) within the observed population (Figure 1). In minimal medium with glucose as a carbon source it was observed that the majority of fluorescent cells showed distinctive foci at the cell poles (Figure 1A). This phenomenon was less obvious following growth in rich medium when GFP fluorescence appeared dispersed throughout the cell (Figure 1B).

**Figure 1:**
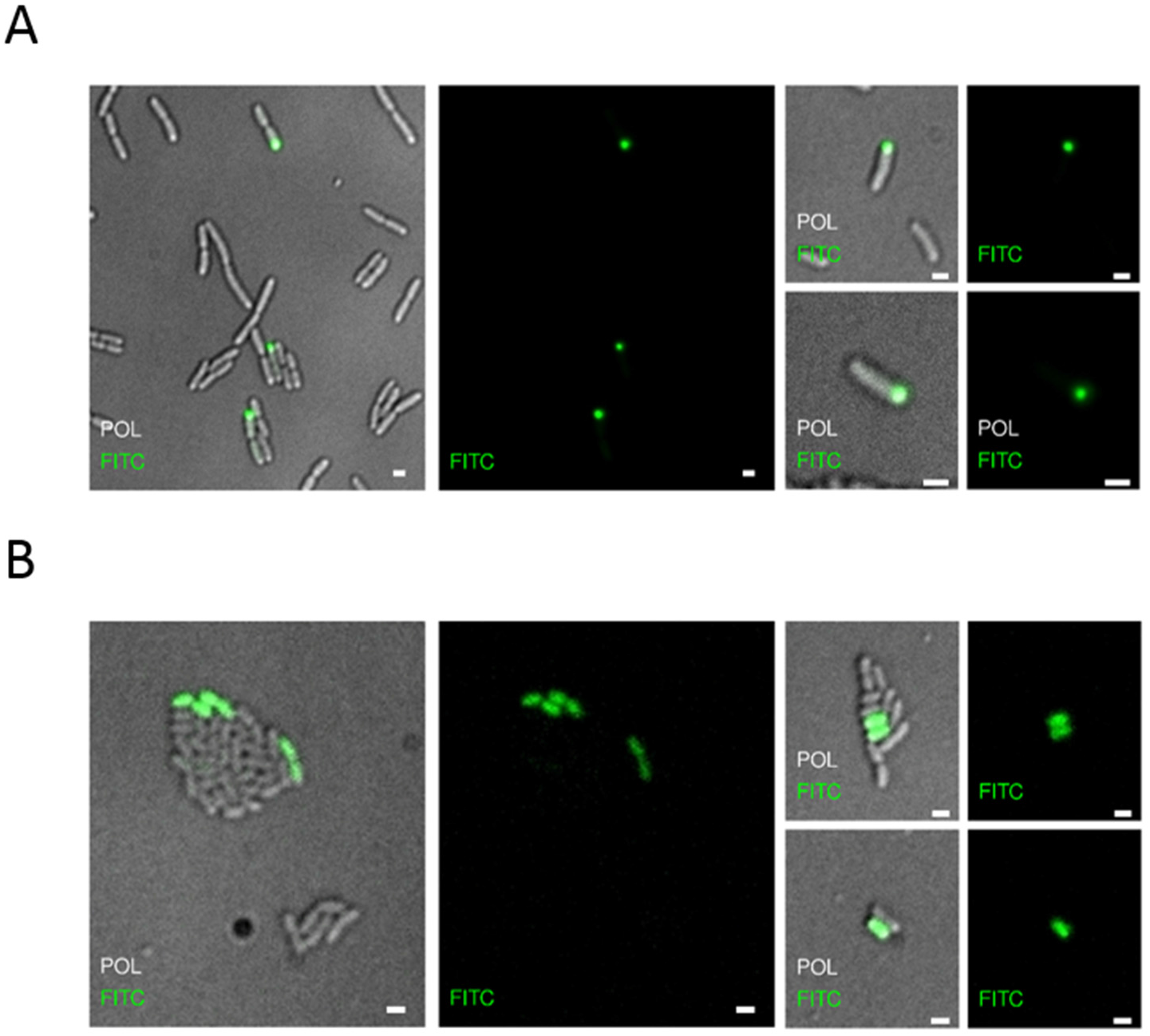
Bimodal expression of ϕ*gfp-chiC*. Representative images of MC03 (DB10, ϕ*gfp::chiC*) cells by fluorescent microscopy. **A.** MC03 cells were grown for 16 hours on MM-Glucose prior to mounting on slides for image acquisition. **B** Cells were grown in rich media for 16 hours prior to microscopic observations. Microscopy slides were prepared with 1% (w/v) agarose and cells were washed and diluted with TSB prior to mounting. Images were acquired using a delta vision microscope, CoolSnap camera, 100X objective lens, using Differential interference contrast (DIC) and fluorescence filters. FITC filter (excitation 490/20; emission 525/30) for GFP detection. Post-acquisition analysis were done on OMERO software. Scale bars of 1 μm.

**Table 1:**
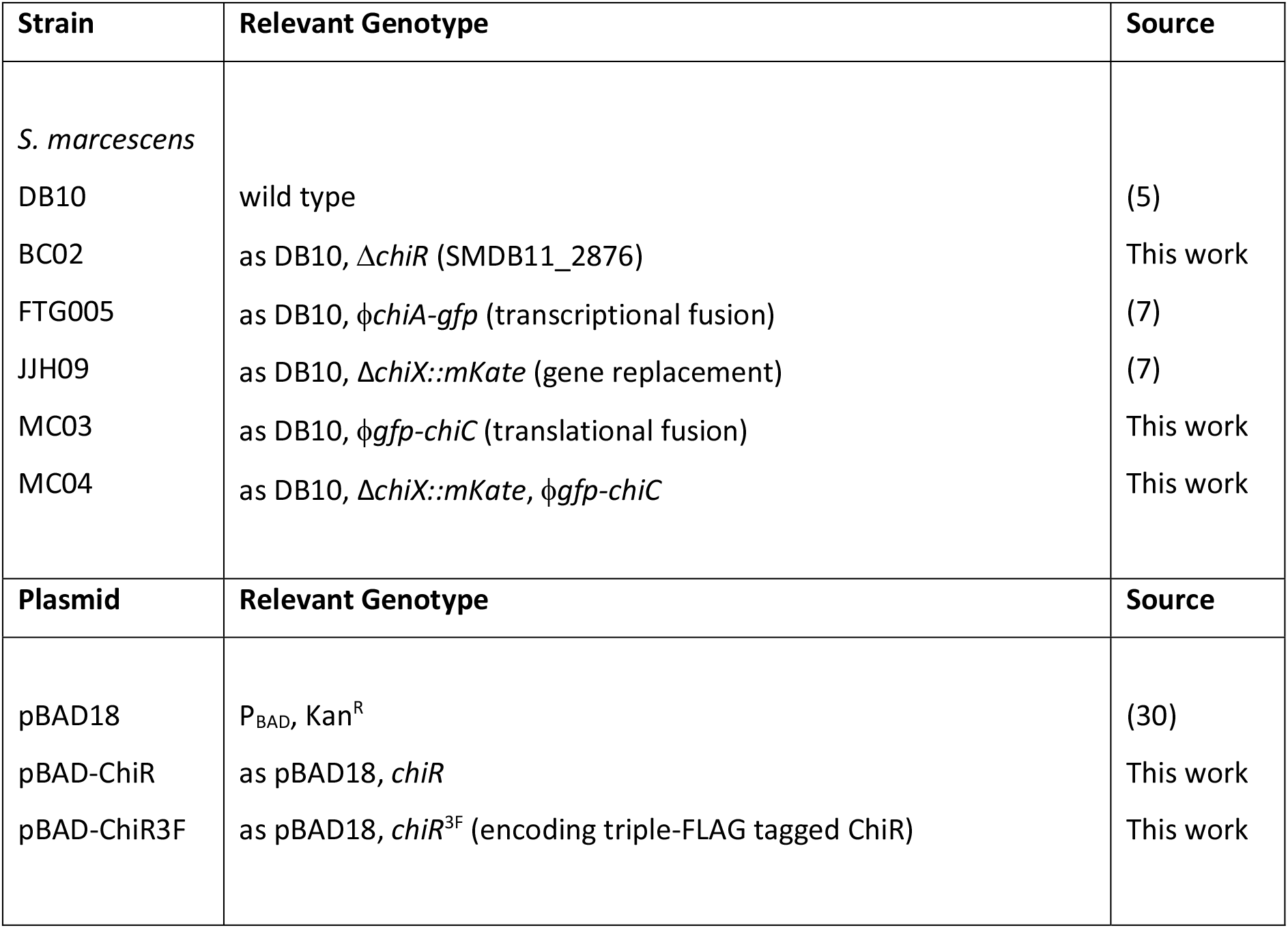
Bacterial strains and plasmids central to this study.

| Strain | Relevant Genotype | Source |
| --- | --- | --- |
| *S. marcescens* |  |  |
| DB10 | wild type | (5) |
| BC02 | as DB10, $\Delta chiR$ (SMDB11_2876) | This work |
| FTG005 | as DB10, $\phi chiA-gfp$ (transcriptional fusion) | (7) |
| JJH09 | as DB10, $\Delta chiX::mKate$ (gene replacement) | (7) |
| MC03 | as DB10, $\phi gfp-chiC$ (translational fusion) | This work |
| MC04 | as DB10, $\Delta chiX::mKate$, $\phi gfp-chiC$ | This work |
| Plasmid | Relevant Genotype | Source |
| pBAD18 | P<sub>BAD</sub>, Kan<sup>R</sup> | (30) |
| pBAD-ChiR | as pBAD18, *chiR* | This work |
| pBAD-ChiR3F | as pBAD18, *chiR*<sup>3F</sup> (encoding triple-FLAG tagged ChiR) | This work |

Flow cytometry was used to quantify the number of ‘ON’ fluorescent cells in the population, with the wild-type *S. marcescens* DB10 strain used as a negative control. MC03 (ϕ*gfp-chiC*) cells were grown in minimal medium containing glucose (MM-Glucose) for 14 hours at 30 °C before being washed in PBS and then screened by flow cytometry (Figure 2A). Data from 10,000 events revealed that the GFP-ChiC fusion was produced in 7.7% of the population (Figure 2A). *S. marcescens* is able to utilise a number of alternative carbon sources and these were also tested here. For instance, growth of the cells in minimal medium containing fructose (MM-Fructose) resulted in 1.4% of the population being ‘ON’ for ϕ*gfp-chiC* expression (Figure 2B). However, again using flow cytometry to facilitate quantification, a slight increase in relative ‘ON’ population was observed when MC03 was grown in glycerol as sole carbon source (15%) or in GlcNAc (13%), which is the basic building block of chitin (Supp. Figure S1).

**Figure 2:**
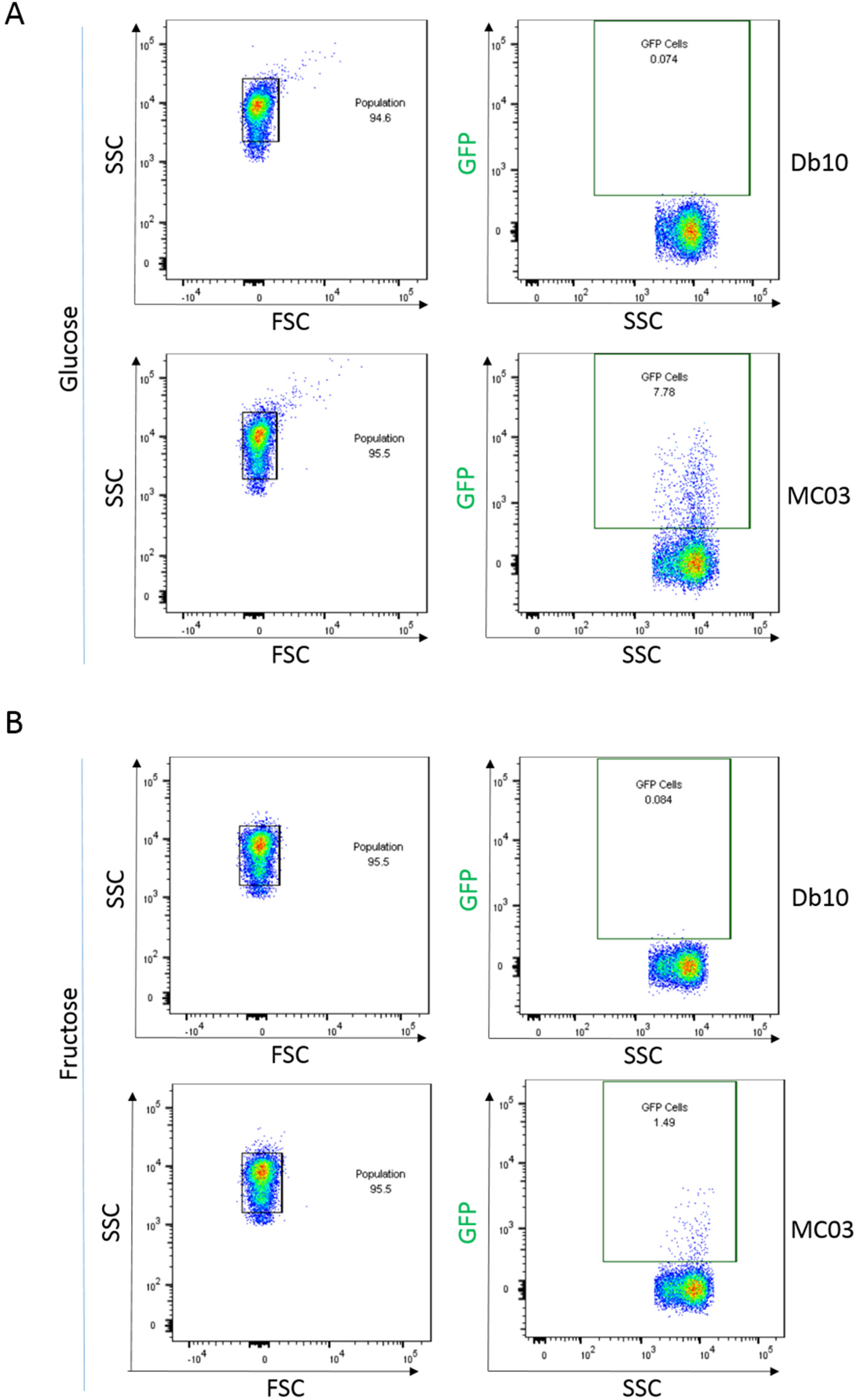
Quantification of bimodal expression of ϕ*gfp-chiC* by flow cytometry. Flow cytometric analysis compared MC03 to the DB10 control. Samples were analysed on a LSR Fortessa (Becton Dickinson). Single bacterial cells were identified on the basis of forward scatter (FSC) and side scatter (SSC) and the GFP fluorescence quantified (488 nm excitation, 530 ± 30 nm emission). The non-GFP expressing DB10 control was used to evaluate background fluorescence and GFP positive cells identified on this basis. **A.** Cell growth in MM-Glucose shows a 7.78% of green fluorescent cells. **B.** Cell growth in MM-Fructose shows green fluorescent subpopulation of 1.49%. Image analysis and figures made using FlowJo v.10.1r7.

Taken together, these data establish that *chiC* expression occurs in a bimodal manner with only a small sub-population of cells producing ChiC at any one time, and the size of that sub-population can vary slightly depending on the growth conditions.

### Evidence for co-ordinated production of ChiC with its secretion system

Previous studies using live cell imaging approaches suggested *chiA* and *chiX* expression was tightly coordinated (7). Here, the relationship between *chiC* expression and *chiX* was explored. Initially, the MC03 (ϕ*gfp-chiC*) strain was modified by the incorporation of a Δ*chiX-mKate* allele to yield strain MC04 (as DB10, Δ*chiX-mKate*, ϕ*gfp-chiC*, Table 1). Next, the double-tagged strain was grown in rich medium for 16 hours, diluted and analysed by fluorescence microscopy (Figure 3). The resultant images clearly showed a small subpopulation of fluorescent cells and in most cases these cells displayed both red and green fluorescence (Figure 3), demonstrating co-ordinated expression of *chiC* and *chiX*.

**Figure 3:**
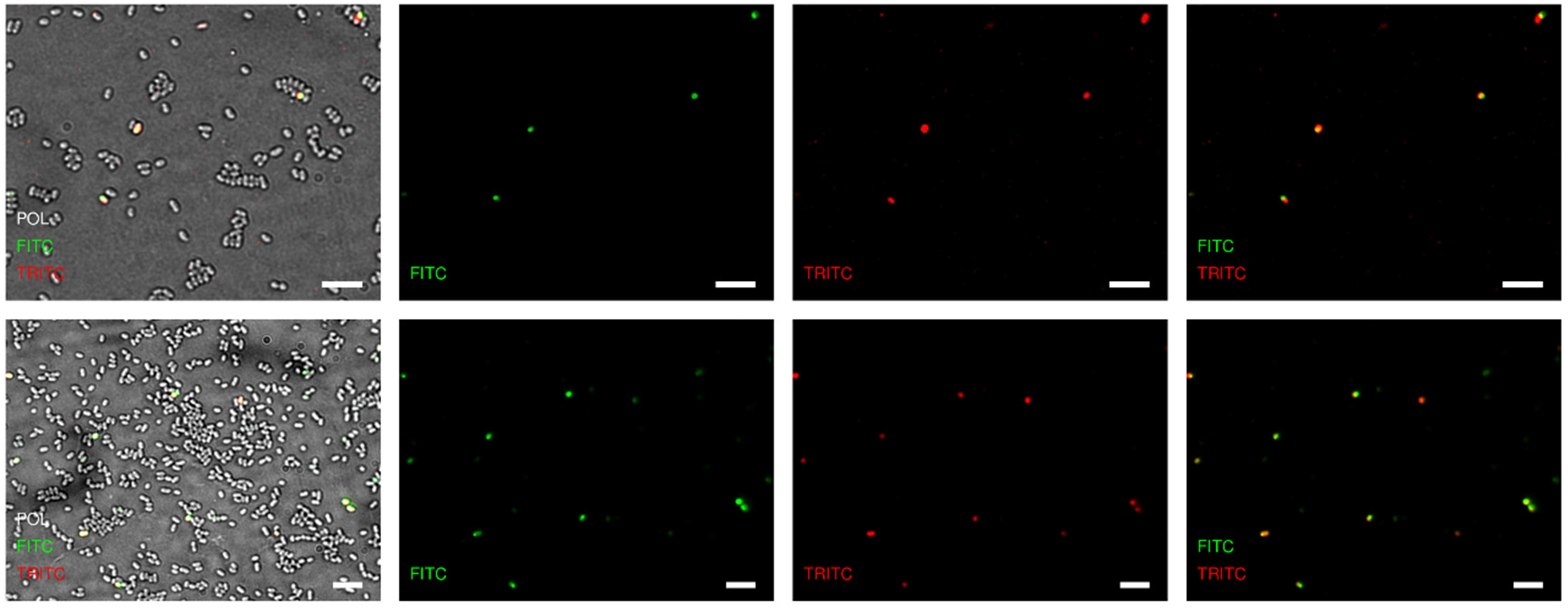
Coordination of *chiC* and *chiX* expression as observed by fluorescence microscopy. Representative images of MC04 (Δ*chiX-mKate*, ϕ*gfp-chiC*) cells containing GFP and mKate as fluorescent reporters of *chiC* and *chiX*, respectively. Cells were grown for 16 hours in rich media prior to mounting on slides for image acquisition. Microscopy slides were prepared with a 1% (w/v) agarose and cells were washed and diluted with TSB prior to mounting. Images were acquired on a Delta Vision microscope, CoolSnap camera, and 100 × objective lens using Differential Interference Contrast (DIC) and fluorescence. FITC filter (excitation 490/20; emission 525/30) was used for GFP detection and a TRITC (tetramethylrhodamine) filter (excitation 555/28; emission 617/73) for mKate detection. Post-acquisition analysis was done on OMERO software. Scale bars, 5 μm.

### Overproduction of ChiR shifts the bimodal expression ratios of *chiA, chiC* and *chiX*

A candidate for controlling the regulation of chitinase production and secretion is ChiR (7, 9, 11). To explore this experimentally, a strain was constructed in which *chiR* alone was deleted (BC02, as DB10 Δ*chiR*, Table 1). The BC02 (Δ*chiR*) strain was grown in rich media and then separated into whole cells and culture supernatant fractions to assess chitinase synthesis and secretion by Western immunoblotting (Figure 4). Neither ChiA nor ChiC proteins were detectable neither inside nor outside the cell in the absence of *chiR* (Figure 4). Complementation experiments were carried out *in trans* using the *S. marcescens chiR* gene encoded on the L-arabinose-inducible pBAD18 (Kan^R^) vector (Table 1). After 14 hours of growth in rich media, BC02/pBAD-ChiR cells were supplemented with 0.005% (w/v) arabinose (final concentration) for a further five hours before cultures were fractionated into whole cell and supernatant samples. Western immunoblotting showed chitinase production and secretion was restored when *chiR* was supplied on a plasmid (Figure 4).

**Figure 4:**
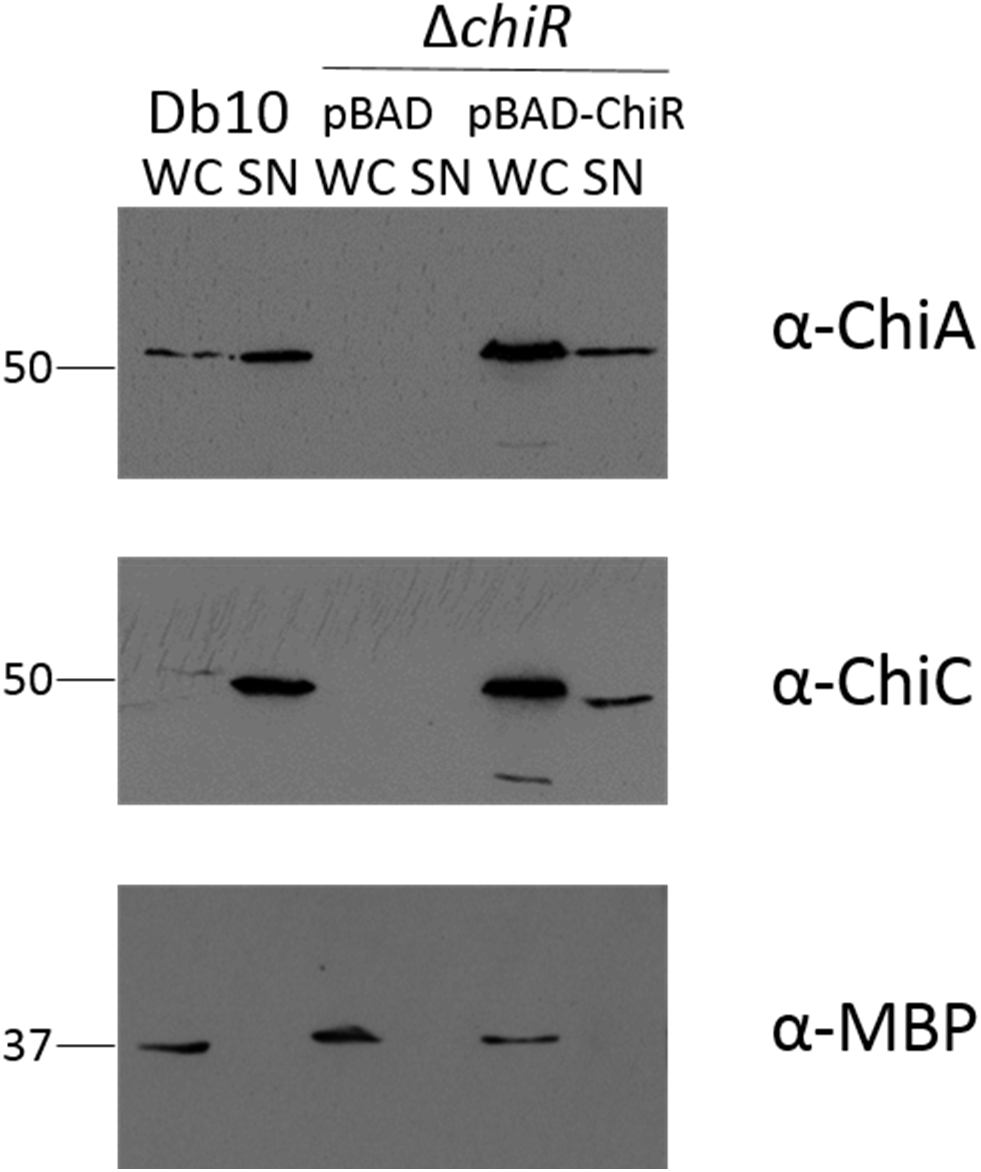
ChiR is essential for chitinase production and secretion. *S. marcescens* wild type DB10 and BC02 (Δ*chiR*) cells transformed with pBAD18 or pBAD-ChiR vector were grown for 14 hours in rich medium before addition of arabinose (0.005% w/v final concentration) for a further 5 hours. Cells were separated into whole cells (WC) and supernatant (SN) before being analysed by SDS-PAGE and Western blotting. Immunoblotting was carried out using the antibodies shown. Maltose binding protein (MBP) was used as a periplasmic (cellular) control.

Since the ChiA and ChiC proteins were no longer detected in the absence of *chiR*, a reverse transcriptase PCR (RT-PCR) experiment was performed to assess transcription of the genes encoding all three chitinases and the *chiWXYZ* operon (Supp. Figure S2). The data revealed that no mRNA transcripts were detectible for *chiA, chiB, chiC, chiW or chiX* in the Δ*chiR* strain (Supp. Figure S2). These results confirm that ChiR is a member of a pathway that transcriptionally regulates the expression of these genes.

Next, the fluorescent reporter strains FTG005 (ϕ*chiA::gfp*), MC03 (ϕ*gfp::chiC*) and JJH09 (Δ*chiX::mKate*) were transformed with pBAD-ChiR3F, which encodes ChiR carrying a Triple-Flag-Tag at its C-terminus. Cells were grown in rich media for 14 hours at 30 °C before L-arabinose was added to the culture at a final concentration of 0.005% (w/v) for 5 hours. Fluorescence microscopy was then applied to assess the production of these fusion proteins in the populations of cells. The images showed clear changes to the bimodal expression patterns of these fusions and that inclusion of the *chiR* plasmid strikingly increased the quantity of fluorescent cells observed in all cases (Figure 5). Furthermore, the co-ordinated expression of the *chiX* gene with *chiC* was found to be preserved when extra ChiR was supplied in the double-fluorscent reporter strain MC04 (ϕ*gfp::chiC*, Δ*chiX::mKate*) (Figure 6).

**Figure 5:**
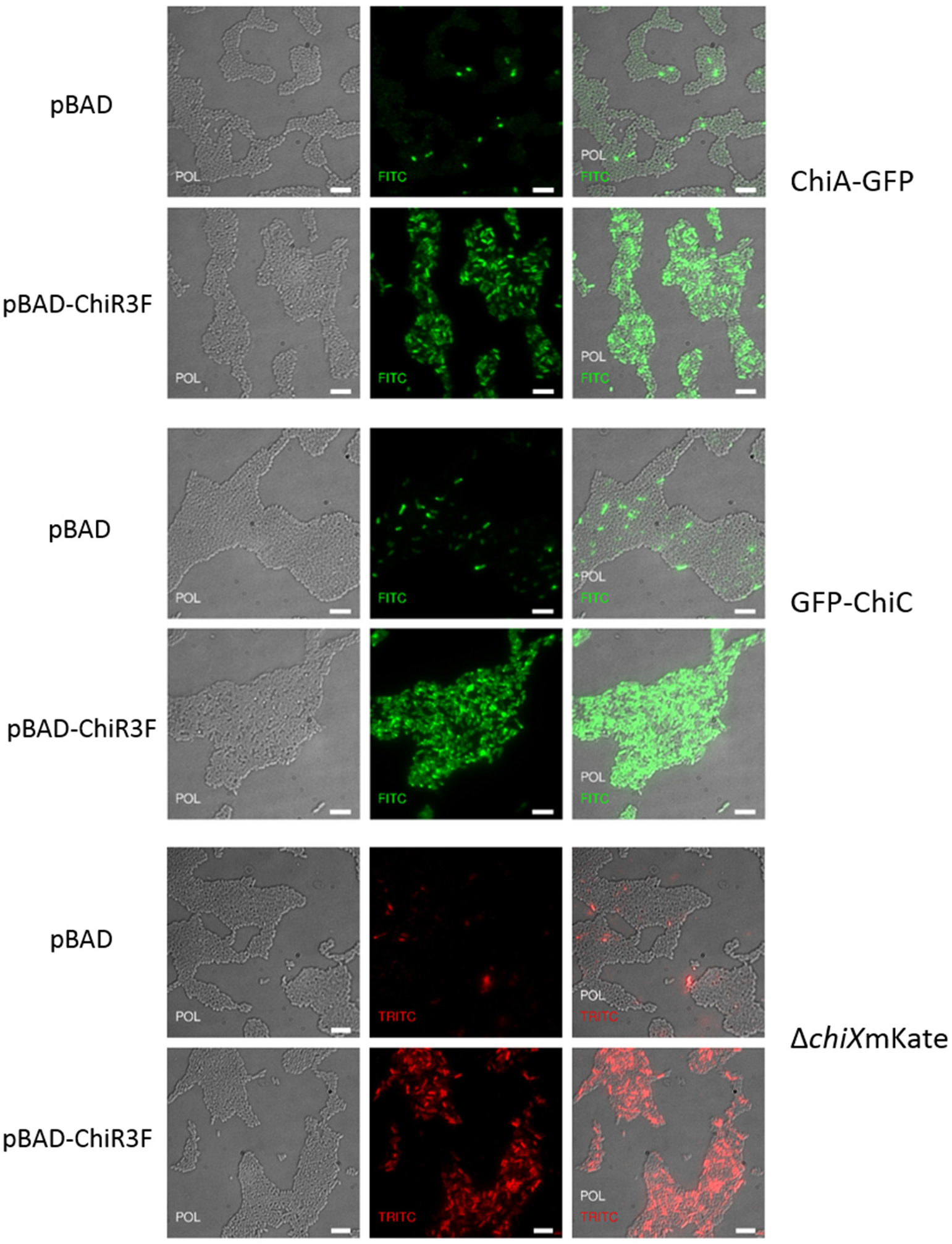
Overexpression of *chiR* increases the population of cells producing chitinases. Representative images of the transcriptional ϕ*chiA-gfp* fusion strain (FTG005), the translational ϕGFP-ChiC strain (MC03) and the ϕ*chiX::mKate* strain (JJH09). The strains were transformed with pBAD control plasmids or pBAD-ChiR3F. Cells were grown for 14 hours in rich medium prior to plasmid expression induction for 5 hours in presence of arabinose (0.005% w/v final concentration). Microscopy slides were prepared in 1% (w/v) agarose and cells were washed and diluted in TSB before mounting. Images were acquired on a Delta Vision microscope using a CoolSnap camera and 100 × objective lens using Differential Interference Contrast (DIC) and fluorescence. Fluorescent filters; FITC filter (excitation 490/20; emission 525/30) and TRITC filter (excitation 555/28; emission 617/73). Post-acquisition analysis was done on OMERO software. Scale bars, 5 μm.

**Figure 6:**
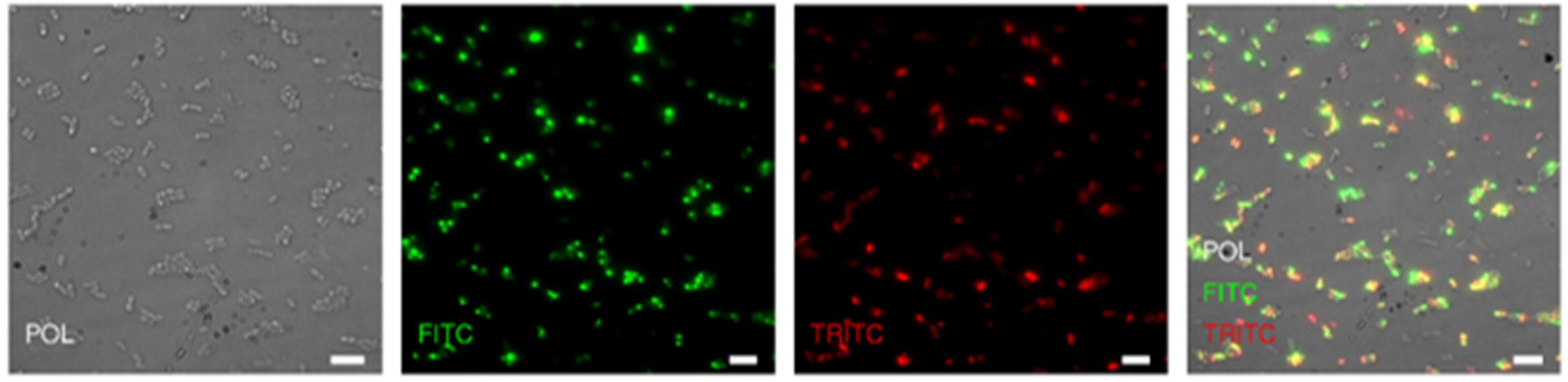
Overexpression of *chiR* preserves coordination between *chiX* and *chiC* genes. Representative image of MC04 (ϕ*gfp::chiC*, Δ*chiX::mKate*) cells under overexpression of *chiR* (pBAD-ChiR3F). Cells were grown in rich medium for 14 hours and induced for expression (0.005% w/v L-arabinose final concentration) for the following 5 hours. Images were acquired on a delta vision microscope, CoolSnap camera, and 100 × objective lens using Differential Interference Contrast (DIC) and filters FITC for GFP detection, and TRITC for mKate detection. Post-acquisition analysis was done on OMERO software. Scale bars, 5 μm.

In order to quantify the observed ChiR effect on *chiC* expression, flow cytometry was performed. Note that due to absence of a laser within the excitation spectrum of mKate, and the failure to consistently detect GFP fluorescence in the cytometer when using the *chiA* fusion strain, all further experiments were performed using the MC03 strain that produces the suitably bright GFP-ChiC fusion. The MC03 (ϕ*gfp::chiC*) cells were transformed with pBAD-ChiR3F and grown in MM-fructose for 14 hours after which L-arabinose was added for 5 hours before the cells were sorted. Cytometry results revealed that under *chiR* overexpression conditions some cells (calculated at 23% of the population tested) displayed an unusual lower side scattering (SSC) (Figure 7). Data from this low SSC sub-population was analysed separately from the remainder of the cells and only 7.2% of these cells could be considered ‘ON’ for GFP fluorescence (Figure 7). However, under ChiR overproduction conditions, the sub-population of cells with more usual SSC were observed to exhibit 79.2% ‘ON’ for GFP fluorescence (Figure 7). Overall, this suggests an increase from 1% to 62% ‘ON’ for GFP fluorescence in the entire population of cells simply by introduction of a ChiR-encoding plasmid.

**Figure 7:**
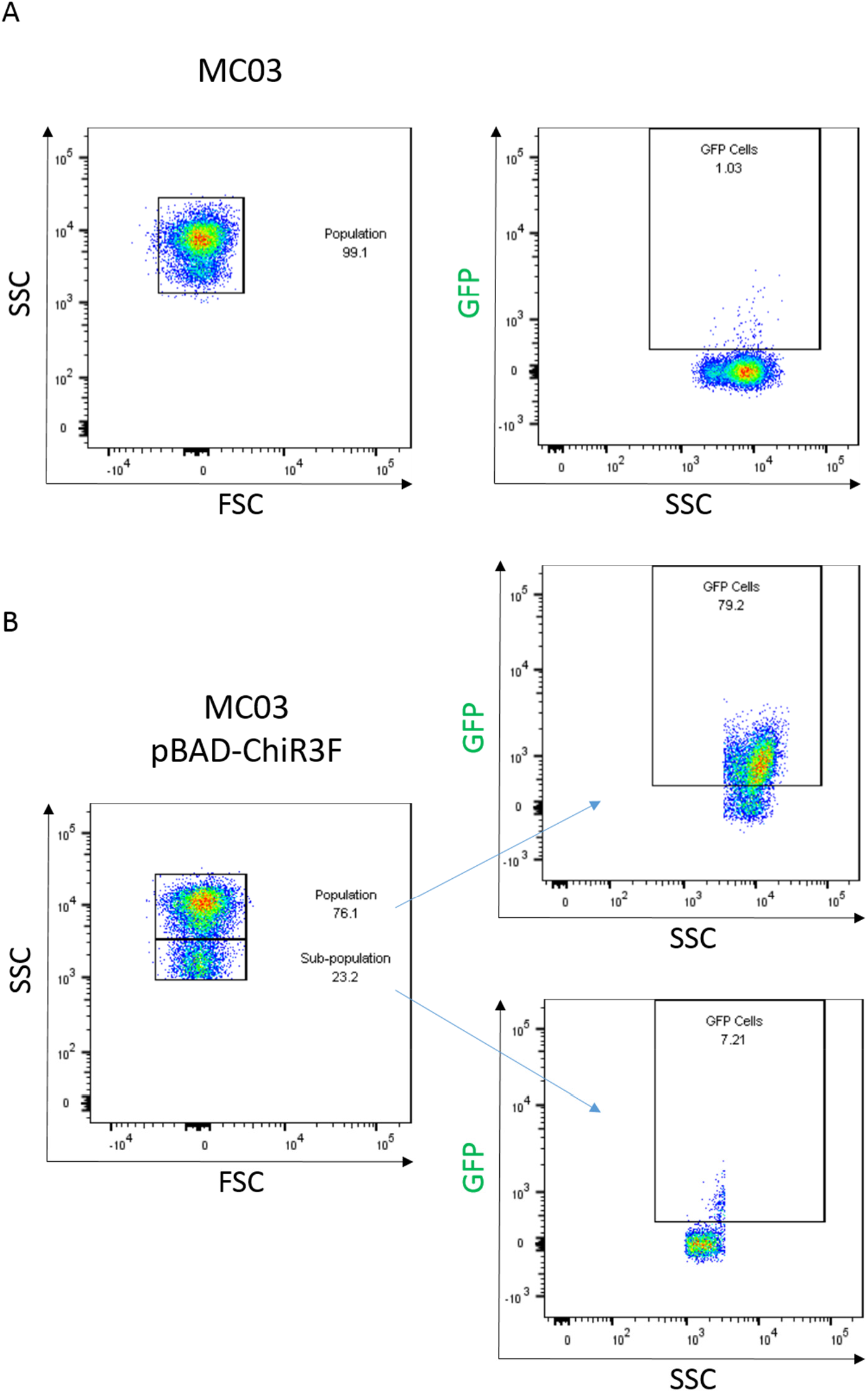
Quantification of the shift in bimodal expression ratio by ϕ*gfp-chiC* induced by ChiR overproduction. Flow cytometric analysis was used to quantify MC03 cells transformed with pBAD-ChiR3F. Cells were grown for 14 hours in minimal medium with fructose prior to induction for 5 hours by addition of arabinose (0.005% w/v final concentration). Samples were analysed on a LSR Fortessa (Becton Dickinson) in which single bacterial cells were identified on the basis of forward scatter (FSC) and side scatter (SSC) and GFP fluorescence. FITC filter (488 nm excitation, 530 ± 30 nm emission). Image analysis and figures were prepared using FlowJo v.10.1r7 software.

Taken together, these data demonstrate that ChiR is important for transcription of *chiA, chiC* and *chiX*, and that increased expression of *chiR* alone is sufficient to significantly shift the physiology of the population from one where chitinase production is rare, to one where chitinase production is operational in the majority of cells.

### A wider influence of ChiR on bacterial physiology revealed by quantitative proteomics

Given the behaviour of *chiA, chiC* and *chiX* in response to ChiR overproduction, it was considered that ChiR activity could be intimately linked with the production of all proteins involved in chitin metabolism in *S. marcescens*. Moreover, the ability of excess ChiR to activate chitinase expression and secretion in a majority of the population meant that quantitative proteomics could be an ideal approach to gain a global view of the wider influence of ChiR on the physiology of the bacterium.

For the proteomics experiments a label-free quantitative mass spectrometry approach was taken with a focus on the relative levels of cellular proteins, rather than the extracellular secretome. The parental strain *S. marcescens* DB10 containing the pBAD18 empty vector, and the BC02 (Δ*chiR*) strain containing either pBAD18 or pBAD-ChiR3F, were utilised for the whole cell proteomic experiments. In each case, four biological replicates were grown in rich media for 14 hours at 30 °C, before supplementation with 0.2% (w/v) L-arabinose for a further 5 hours. Cultures were then harvested and washed in PBS buffer before proteins were extracted, tryptic peptides prepared, and LC-MS/MS analysis carried out. All samples were highly correlated (R^2^ > 0.9) and Supp. Table S1 shows the number of unique peptides and proteins detected in each replicate.

### Comparison of the *S. marcescens* DB10 proteome with that of *ΔchiR* strain

First, a comparison of the proteome of a strain with native-level ChiR activity, using the DB10 wild-type peptide data, was compared with the proteome of the BC02 (Δ*chiR*) strain. The comparison is represented as a volcano plot (Figure 8) and in Table 2. In this case, proteins significantly more abundant in DB10 (log2 >2.0, p <0.05) were ChiA, Cbp21 and ChiC (Table 2). The ChiB protein was also detected in the data set, but its relative abundance (log2 +1.7) fell below the log2 >2.0 threshold employed here (a comprehensive data comparison is provided in Supp. Table S2).

**Figure 8:**
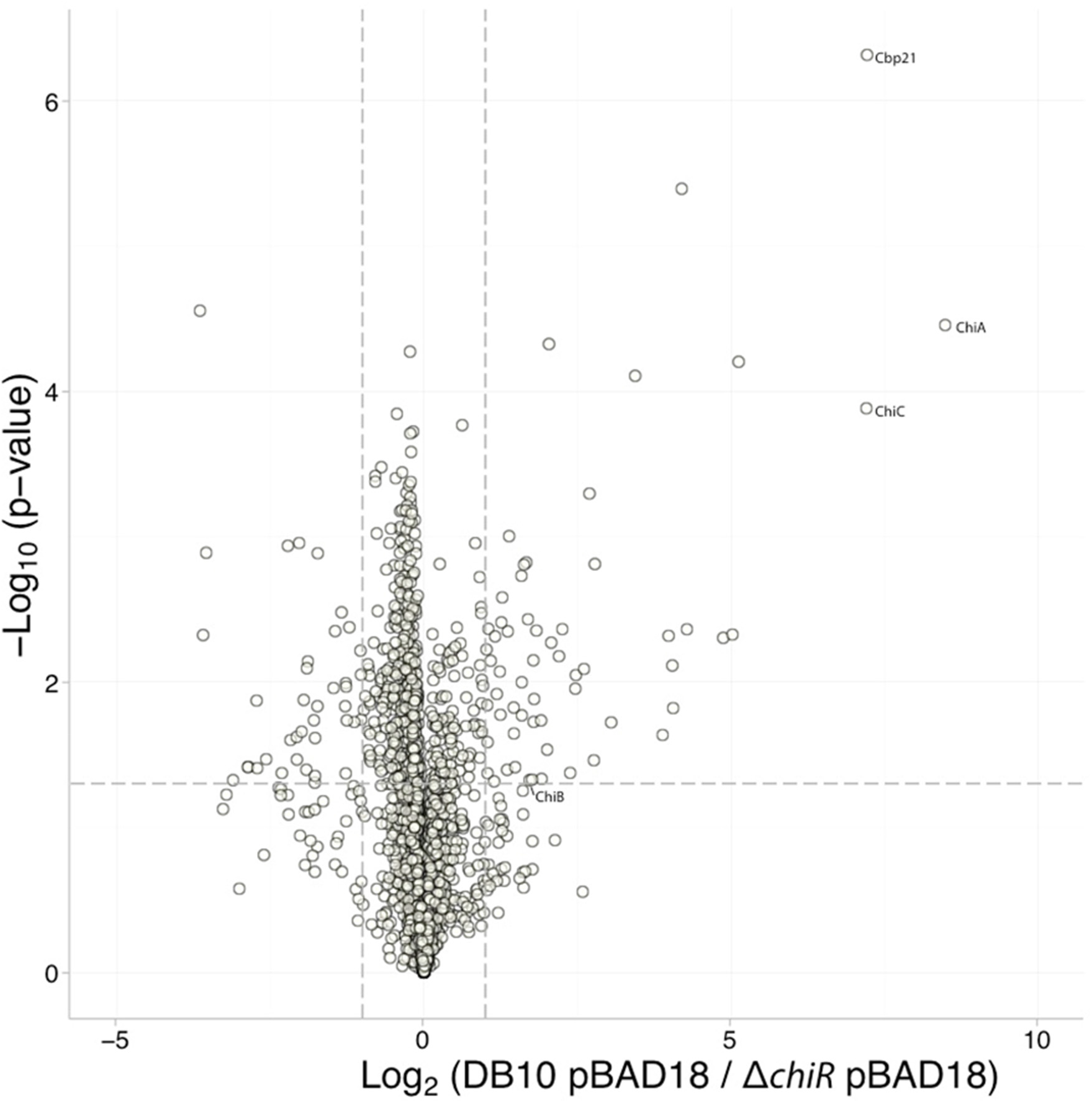
Comparison of the cellular proteome of *S. marcescens* DB10 with a Δ*chiR* mutant. A volcano plot demonstrates the changes in relative abundance of proteins identified in *S. marcescens* DB10 carrying empty vector pBAD18 *versus* BC02 (Δ*chiR*) carrying empty vector pBAD18. The locations of the chitinolytic machinery in the plot is indicated.

**Table 2:**
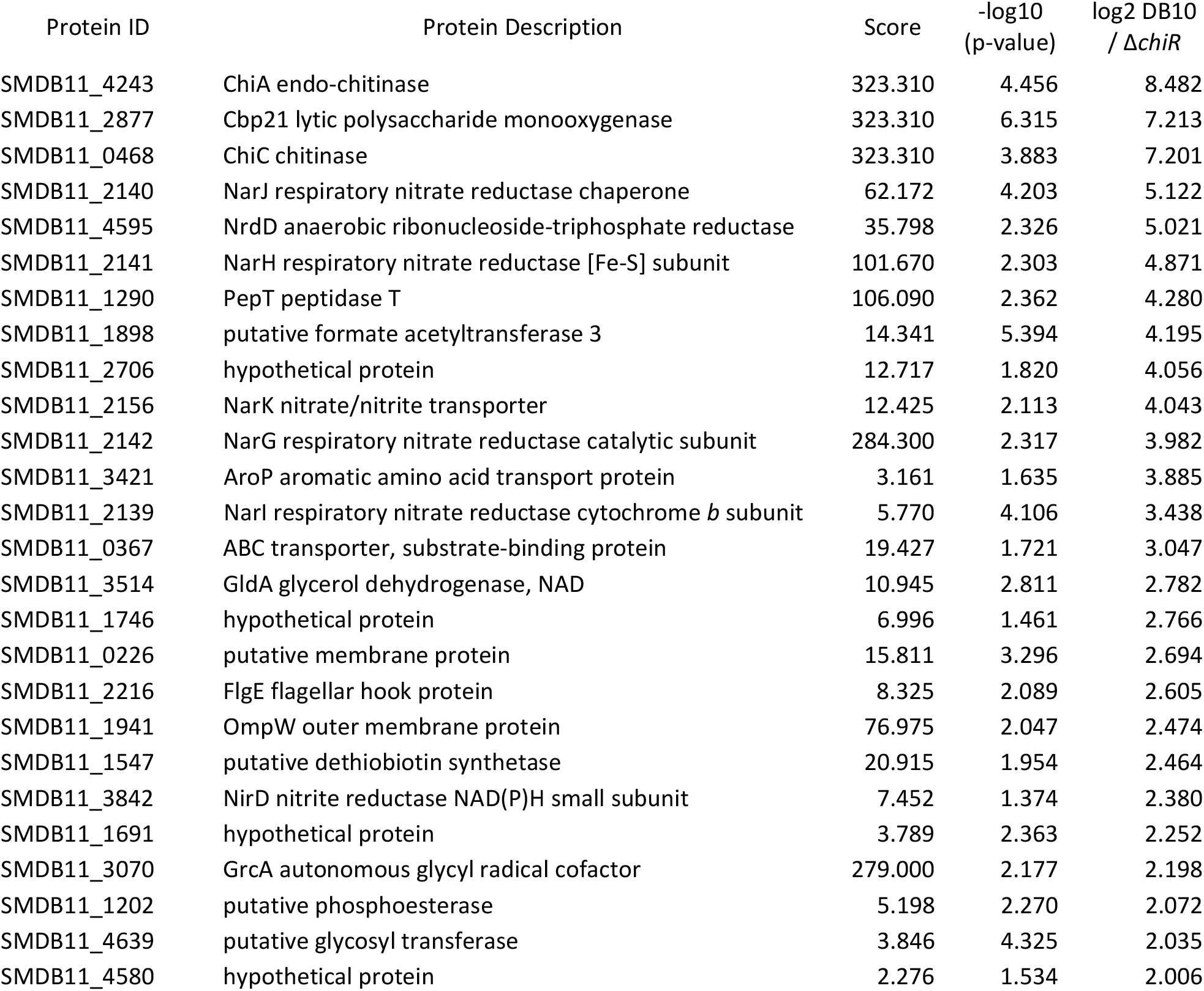
Proteins more abundant in the *S. marcescens* BD10 wild-type strain compared to a Δ*chiR* mutant. Proteins showing a significant change (log2 >2.0, p <0.05) in relative abundance are shown alongside their Andromeda score. Four biological replicates of each strain were analysed.

A surprising feature of these data was the identification of the respiratory nitrate reductase proteins being influenced by cellular ChiR levels in *S. marcescens* (Table 2). The NarGHI complex is a membrane-bound metalloenzyme that links quinol oxidation to nitrate reduction. All three nitrate reductase subunits, plus a biosynthetic chaperone NarJ encoded within the same operon, were identified as being significantly negatively affected upon deletion of *chiR* (Table 2). Strikingly, other players in nitrogen metabolism, such as the membrane transporter NarK and the cytoplasmic NAD(P)H-dependent nitrite reductase, were also notably negatively affected by the absence of ChiR (Table 2).

### Comparison of the *S. marcescens* ΔchiR proteome with that overproducing ChiR

Next, a comparison was made between samples from the BC02 (Δ*chiR*) strain producing extra ChiR from a plasmid and those carrying only the empty vector pBAD18. The comparison is represented as a volcano plot (Figure 9) and in Table 3. Proteins significantly more abundant in BC02 (Δ*chiR*) when complemented with pBAD-ChiR3F (log2 >2.0, p <0.05) were ChiA, Cbp21, ChiC and ChiB (Table 3). In addition, products of the chitinase secretion operon (ChiX and ChiY) were detected and shown to be significantly more abundant in cells overproducing ChiR (Table 3). The respiratory nitrate reductase and the NarK transporter are also prominent as the most severely affected proteins upon overexpression of *chiR* (Table 3). A comprehensive data comparison is provided in Supp. Table S3.

**Figure 9:**
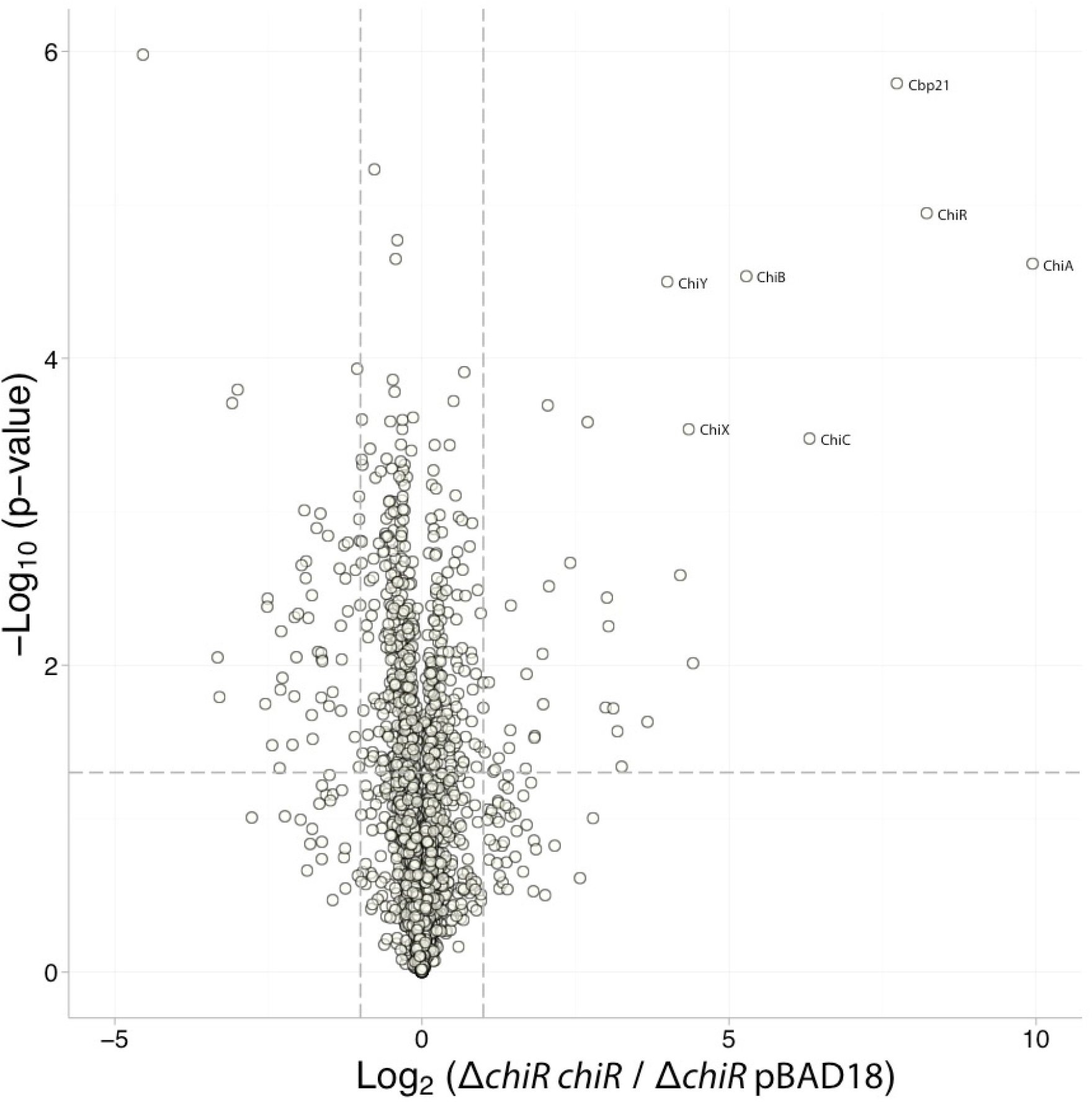
Comparison of the cellular proteome of an *S. marcescens Δ*chiR** mutant with that of a complemented mutant overproducing ChiR. A volcano plot demonstrates the changes in relative abundance of proteins identified in *S. marcescens* BC02 (Δ*chiR*) carrying vector pBAD18-ChiRTF encoding extra ChiR *versus* BC02 (Δ*chiR*) carrying empty vector pBAD18. The locations of the chitinolytic machinery in the plot is indicated.

**Table 3:**
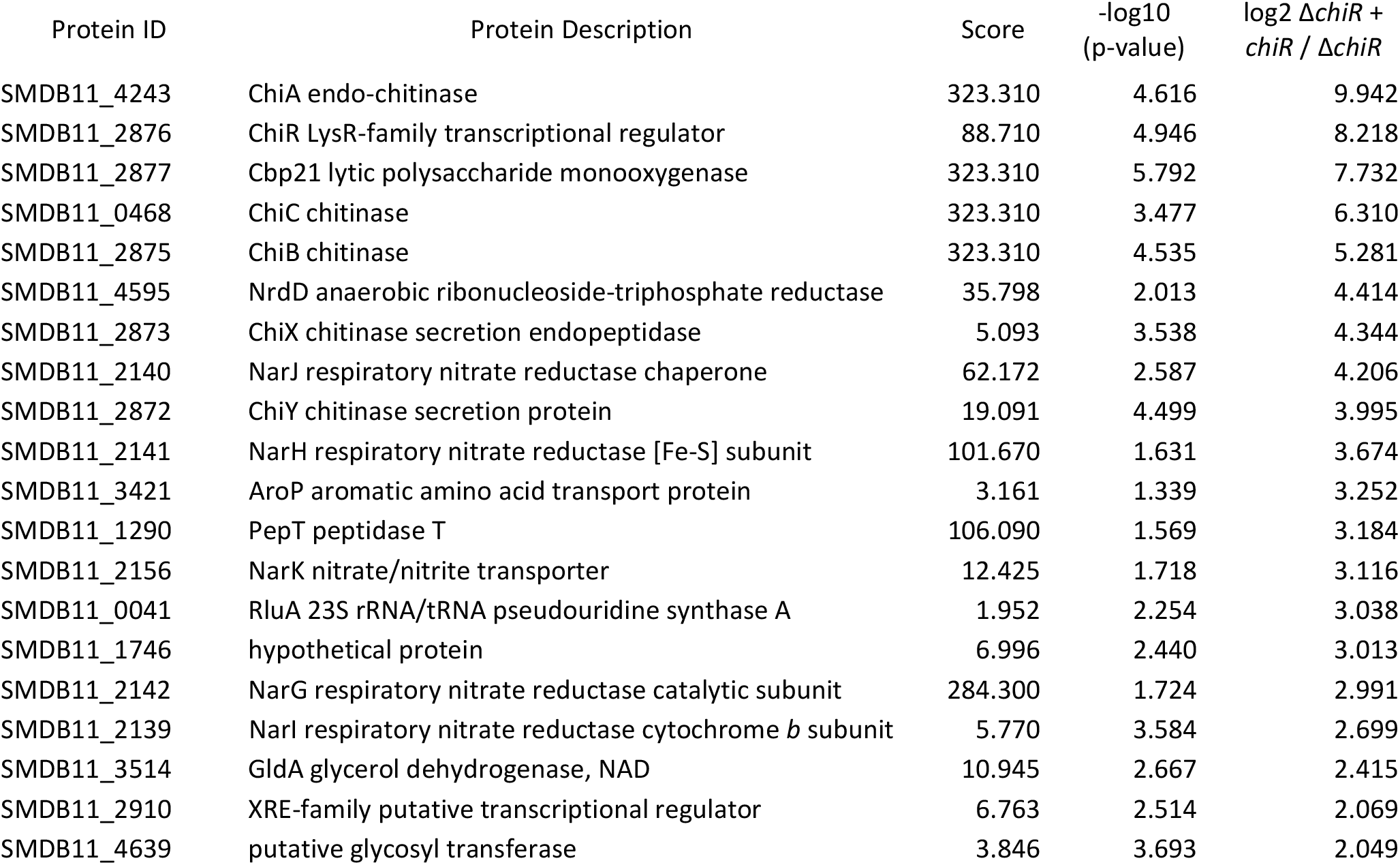
Proteins more abundant in the *S. marcescens* Δ*chiR* mutant overexpressing *chiR* compared to the Δ*chiR* mutant alone. Proteins showing a significant change (log2 >2.0, p <0.05) in relative abundance are shown alongside their Andromeda score. Four biological replicates of each strain were analysed. No peptides from ChiR were detected in the Δ*chiR* mutant, the ratio was determined by imputing correlating intensities around the detection limit.

| Protein ID | Protein Description | Score | $-\log_{10}$<br>(p-value) | $\log_2 \Delta chiR +$<br>$chiR / \Delta chiR$ |
| --- | --- | --- | --- | --- |
| SMDB11_4243 | ChiA endo-chitinase | 323.310 | 4.616 | 9.942 |
| SMDB11_2876 | ChiR LysR-family transcriptional regulator | 88.710 | 4.946 | 8.218 |
| SMDB11_2877 | Cbp21 lytic polysaccharide monooxygenase | 323.310 | 5.792 | 7.732 |
| SMDB11_0468 | ChiC chitinase | 323.310 | 3.477 | 6.310 |
| SMDB11_2875 | ChiB chitinase | 323.310 | 4.535 | 5.281 |
| SMDB11_4595 | NrdD anaerobic ribonucleoside-triphosphate reductase | 35.798 | 2.013 | 4.414 |
| SMDB11_2873 | ChiX chitinase secretion endopeptidase | 5.093 | 3.538 | 4.344 |
| SMDB11_2140 | NarJ respiratory nitrate reductase chaperone | 62.172 | 2.587 | 4.206 |
| SMDB11_2872 | ChiY chitinase secretion protein | 19.091 | 4.499 | 3.995 |
| SMDB11_2141 | NarH respiratory nitrate reductase [Fe-S] subunit | 101.670 | 1.631 | 3.674 |
| SMDB11_3421 | AroP aromatic amino acid transport protein | 3.161 | 1.339 | 3.252 |
| SMDB11_1290 | PepT peptidase T | 106.090 | 1.569 | 3.184 |
| SMDB11_2156 | NarK nitrate/nitrite transporter | 12.425 | 1.718 | 3.116 |
| SMDB11_0041 | RluA 23S rRNA/tRNA pseudouridine synthase A | 1.952 | 2.254 | 3.038 |
| SMDB11_1746 | hypothetical protein | 6.996 | 2.440 | 3.013 |
| SMDB11_2142 | NarG respiratory nitrate reductase catalytic subunit | 284.300 | 1.724 | 2.991 |
| SMDB11_2139 | NarI respiratory nitrate reductase cytochrome *b* subunit | 5.770 | 3.584 | 2.699 |
| SMDB11_3514 | GldA glycerol dehydrogenase, NAD | 10.945 | 2.667 | 2.415 |
| SMDB11_2910 | XRE-family putative transcriptional regulator | 6.763 | 2.514 | 2.069 |
| SMDB11_4639 | putative glycosyl transferase | 3.846 | 3.693 | 2.049 |

## DISCUSSION

### ChiR controls bimodality of expression of the chitinolytic machinery

Bimodal gene expression is an important aspect of bacterial physiology and underpins the molecular basis of phenotypic variability in otherwise ‘pure’ cultures of bacteria. The commitment of only a small sub-population of a culture to a specific type of metabolism has been described as a type of ‘biological bet hedging’ for the wider population (14). In this case of chitin metabolism studied here, the data reveal that 1-14% of the population (slight variations caused by different growth conditions) are ready to contribute to extracellular chitin breakdown at any point in time (Figure 2, Supp. Figure S1). Importantly, the data described here not only show that ChiR important for expression of chitinases *per se* (9), but more interestingly now establish ChiR as the critical player in controlling co-ordinated bimodal chitinase gene expression in *S. marcescens*. Deliberate production of ChiR across the culture by using an plasmid-based system, had the dramatic effect of not simply increasing the amount of chitinase produced by 1-14% of the population, but by increasing the ratio of cells producing and secreting chitinases to 62% (Figure 7). This suggests stochastic fluctuations in cellular ChiR levels may be the sole determinant of chitinase production in *S. marcescens*, though the existence of additional more elaborate regulatory factors cannot be ruled out. Indeed, ChiR was originally identified in genetic screens focused on screening for chitinase activity (7, 9) and was later exploited as a positive regulator of chitinase activity in biotechnological applications (11). Regulation of chitinases is thought to be further influenced by the presence of chitobiose (the GlcNAc disaccharide) (15) and by small regulatory RNAs (16). We have no evidence here that a significant shift in bimodality would occur if a chitin source or breakdown were to be encountered by the population, but since growth on GlcNAc as sole carbon source led to 13% of cells ‘ON’ for GFP-ChiC production, this may be unlikely (Supp. Figure S1).

### Quantitative proteomics reveals cells braced to encounter nitrogen compounds

As well as determining the extent of bimodal gene expression, data described in this work also establishes ChiR for the first time as an important player in the co-ordination of expression of the entire chitinolytic machinery. First, live cell imaging revealed that expression of the *chiWXYZ* secretion operon is perfectly co-ordinated with the expression of *chiC* (Figures 5, 6), and this is entirely consistent coordinated expression between *chiA* and *chiWXYZ* (7). Strikingly, the ability of extra ChiR in the cell to induce expression of ChiC in a major portion of the population (Figure 5) is mirrored perfectly by a concomitant increase in the population of cells expressing the secretion system (Figures 5, 6). Second, this tight relationship between ChiR-dependent expression of *chiA, chiC* and *chiX* suggested that other proteins involved in chitin metabolism might be unearthed by focusing on the role of ChiR in the cell. To this end, as set of quantitative label-free proteomics experiments were conducted (Figures 8, 9). This approach allowed the relative abundance of cellular proteins to be compared between different genetic backgrounds and the fine details of the phenotypic differences to be teased out (Tables 2, 3).

By comparing the cellular proteomes of *S. marcesens* completely devoid of ChiR with that overproducing ChiR (Table 3), the first conclusion that could be firmly established was that ChiR coordinately controls all of the known chitinolytic machinery, including ChiB, Cbp21 and ChiY, which were proteins that had never been examined by fluorescent labelling.

The second major conclusion that could be drawn is that, far from chitinase-producing cells demonstrating some sort of ‘altruistic lysis’ or simply playing a role in chitinase production for the wider population, the ChiR producing cells were readying themselves to metabolise the products of chitin breakdown. Chitin is both a carbon and a nitrogen source for *S. marcescens*. Through the action of the cytoplasmic Nag proteins (which are interestingly apparently not regulated by ChiR), GlcNAc is broken down into equimolar quantities of glucose, acetate and ammonia (17). Although ammonia can be assimilated as an essential nitrogen source, excess ammonia is likely to be toxic to the cell. It is clear from the new proteomics data described here that a full pathway of nitrogen metabolism is coordinately produced alongside the chitinolytic machinery (Tables 2, 3). First, the NADH-dependent nitrite reductase (NirD) is induced by excess ChiR (Table 3) and reduced by removing ChiR (Table 2). NirD is part of an enzyme that would convert excess cytoplasmic ammonia into nitrite and NADH (18). Cytoplasmic nitrite in excess is toxic and the cell would normally look to exchange that for any extracellular nitrate in the environment (19). Indeed, a NirK nitrite/nitrate exchanger was also found to be induced upon overproduction of ChiR (Table 3). Finally, having produced a molecule of NADH and potentially brought a molecule of nitrate into the cytoplasm, the cell has the opportunity to conserve some energy in the form of a respiratory electron transport chain (20). Strikingly, overproduction of ChiR clearly induces production of all four products of the *narGHJI* operon, which encodes a membrane-bound quinone-dependent respiratory nitrate reductase (NarGHI) and its biosynthetic chaperone NarJ. Although most commonly an anaerobic activity, such ammonia and nitrate metabolism can also happen under aerobic conditions (21). Co-production of NirD with NarGHI has been observed in Mycobacterium tuberculosis, but in that case the system biased towards assimilation of nitrate into ammonia for cell growth (22). Taken altogether, it is clear *S. marcescens* is bracing itself to metabolise excess nitrogen compounds at the same time as it breaks down extracellular chitin.

### Concluding remarks

In this work it is revealed that, rather than simply controlling levels of chitinase gene transcription within individual cells, ChiR is central to controlling the biomodal pattern of gene expression across the entire population. Typically, an LTTR-type regulator would be expected to bind a small activator molecule in order to be activated. There is no known activator for ChiR as yet, but these small molecules are frequently associated with central metabolism, quorum sensing, or oxidative stress (10). LTTR-type regulators are being implicated in increasingly important roles in microbial metabolism (23), and it is anticipated that further research into the role of *S. marcescens* ChiR will unearth new components of the chitinase secretions system, extra members of the chitinolytic machinery, and a fresh insight into bacterial physiology.

## MATERIALS AND METHODS

### Bacterial strains and growth conditions

The parental strain used in this study is *S. marcescens* DB10 (5) and derivatives listed in Table 1. Gene replacements and deletions in *S. marcescens* were constructed by allelic exchange using the suicide vector pKNG101 (24) essentially as described previously (25). Briefly, for strain BC02, an in-frame Δ*chiR* allele was assembled and sequenced first on pKS^+^ Bluescript (Amp^R^) and then cloned in to pKNG101 before being transferred to *S. marcescens* DB10 by conjugation. Strain MC03 was constructed by first assembling a ϕ*gfp-chiC* translational fusion on pKS^+^ Bluescript. The GFP *mut2* sequence (minus stop codon) was positioned in-frame with the natural promoter and start codon of *chiC* by PCR. Then ~500 bp of sequence encoding the ChiC N-terminus (minus start codon) was cloned in-frame with *gfp* to generate a ϕ*gfp-chiC* allele. The construct was DNA sequenced before the allele was transferred to the chromosome of *S. marcescens* DB10 *via* pKNG101. Strain MC04 was constructed by transferring the ϕ*gfp-chiC* allele to strain JJH09 (Δ*chiX::mKate* (7)).

*S. marcescens* strains were grown aerobically at 30°C in ‘low salt’ LB medium (10 g/l tryptone, 5 g/l yeast extract, 5 g/l NaCl) or in Minimal Medium (MM) (40 mM K_2_HPO_4_, 15mM KH_2_PO_4_, 0.1% (w/v) (NH_4_)_2_SO_4_, 0.4 mM MgSO_4_) supplemented with carbon sources as indicated.

### Microscopy

Cell cultures were grown according the experimental purpose, typically 16 hours for native levels of expression and 19 hours (14 hours growth plus 5 hours of L-arabinose induction) for *chiR* overexpression experiments, with constant shaking at 30°C. Cells were harvested, washed and diluted 1:10 in 1 × PBS buffer. An agarose pad (1.5%) was prepared on top of a microscopy slide using a Gene frame^®^ (Thermo Scientific, 65 μL coverslips 65 μL, 15 mm × 16 mm internal). Then 1 μl of cells was spotted onto it, left to dry under sterile conditions and sealed with a coverslip (1.5 thickness, VWR). Most of images were acquired using a DeltaVision Core widefield microscope (Applied Precision) mounted on an Olympus IX71 inverted stand. An Olympus 100X 1.4 NA lens with Cascade2 512 EMCCD camera (Photometrics) combined with Differential interference contrast (DIC) and fluorescence optics was used. DIC images were acquired using an LED Transmitted light at 32% intensity and exposure of 25-100 ms. Fluorescent proteins, GFP and mKate, were detected using FITC (fluorescein isothiocyanate) (excitation 490/20; emission 528/38) and TRITC (Tetramethylrhodamine) (excitation 555/28; emission 617/73) filters set. The microscope used for ultra-resolution microscopy was an OMX Blaze system (GE Healthcare) with UPlanSApochromat 63 × 1.42NA, oil immersion objective lens (Olympus) and scientific CMOS camera (PCO AG, Germany). Raw images were processed for channels alignment using the acquisition software of the microscope. All microscopy images were visualized using OMERO software (http://openmicroscopy.org) for postacquisition analysis and image preparation.

### Flow cytometry

Cells were grown in minimal medium at 30°C for 14 hours, for native levels of expression, or for an additional 4 hours following L-arabinose addition for experiments with *chiR* overexpression. Samples were harvested by centrifugation, washed in 1 × PBS buffer and diluted. Samples were transferred to a cytometry tube containing 1 ml of 1 × PBS buffer and analysed on a LSR Fortessa (Becton Dickinson) cytometer. Single cells were identified on the basis of Forward Scatter (FSC) and Side Scatter (SSC), and fluorescence identified using a 488 nm laser excitation combined with a FITC band pass filter (530 ± 30). Data analysis were performed on FlowJo v.10.1r7 version software in which samples were visualized on dot plots and population gates generated in the region of major events. Fluorescent cells were identified upon threshold measurement of wild type cells (background fluorescence).

### Proteomics sample preparation

Relevant strains were transformed with either pBAD18 empty vector or pBAD18 ChiR-TF and plated onto 100 μg/l kanamycin 0.5% (w/v) D-glucose LB agar plates. Four colonies were picked from each plate and cells grown in 5 ml LB cultures supplemented with 100 μg/l kanamycin for 14 hours at 30 °C, before induction with 0.2% (w/v) L-arabinose for 5 hours. Cultures were then harvested by centrifugation, washed four times in cold PBS buffer, flash frozen and stored at −80 °C. For cell lysis, 20 μl 5% (v/v) RapiGest™ surfactant was added to each cell pellet, mixed, and then diluted with 50 mM Tris.HCl pH 8.0 to a final RapiGest™ concentration of 1% (v/v). Next, the protein samples were normalised to contain 20 μg of protein in a volume of 10 μl. TCEP was then added to a final concentration of 1 mM, mixed, and heated at 70 °C for 5 minutes. Samples were allowed to cool to room temperature before treatment with 5 mM iodoacetamide for 20 minutes in the dark, followed by treatment with 10 mM DTT for an additional 20 minutes in the dark. Next, trypsinolysis was performed. Samples were diluted 1:10 with 50 mM Tris.HCl pH 8.0 and vortexed. Trypsin was then added at a 1:100 ratio and the samples incubated for 4 hours at 37 °C. This was followed by another addition of trypsin of the same amount, followed by an overnight incubation at 37 °C. TFA was then added to a final concentration of 1% (w/v) to acidify the samples, followed by a 1 hour incubation at 37 °C.

Samples were then centrifuged at 21,000 × g at room temperature for 30 minutes and the supernatants retained. Reversed phase chromatography using a C18 column was then carried out using half of the supernatant (corresponding to 10 μg of initial protein) and desalted peptides were suspended in 0.1% (w/v) TFA to a final concentration of 0.5 mg/ml

### Proteomics Data Collection and Analysis

LC-MS/MS analysis was carried out by the Proteomics and Mass Spectrometry Facility (University of Dundee). Approximately 1 μg of each peptide sample was separated over a two-hour linear acetonitrile gradient on a 50 cm C18 column, with eluting peptides analysed online using Q Exactive HF mass spectrometer.

Raw MS files were analysed utilising a MaxQuant software package (version 1.5.2.8) (26). MS/MS spectra were searched with an integrated Andromeda search engine (27) against a combined database of *S. marcescens* proteins (containing 4724 sequences) alongside a list of common contaminants. Enzyme specificity was set to hydrolyse peptide bonds C-terminal to Lys and Arg with a maximum of two missed cleavage sites allowed per peptide sequence. Carbamidomethylation of Cys was specified as a fixed modification, with oxidation of Met and acetylation protein N-termini selected as variable modifications. Multiplicity was set to one. The “match-between-runs” feature was enabled to transfer peptide identifications to unsequenced or unidentified spectra by matching their masses and retention times. A MaxLFQ algorithm (integrated into MaxQuant software) was applied for label free quantification, with a minimum LFQ ratio count set to one. The processed data was filtered by posterior error probability to achieve a false discovery rate (FDR) of 1% at peptide-to-spectrum matches (PSM) and protein level. The output protein LFQ values were used for downstream analyses (28). Contaminants, and the matches identified based on a reversed decoy sequence database were excluded from the analysis. Proteins were considered identified if they were present in three out of four biological replicates.

The log-transformed distribution of protein LFQ intensity values resembles a truncated normal distribution. Missing values were assumed to primarily occupy this truncated region, and had values imputed corresponding to the proportion of missing proteins from the total proteins identified. Comparisons were then carried out using Perseus software (29) and R scripting language, with > 2-fold change (t-test p-value < 0.05) in relative abundance considered significant and presented in the results section.

## ACKNOWLEDGEMENTS

This work was funded by the Coordenação de Aperfeiçoamento de Pessoal de Nível Superior (CAPES) PhD program of Brazil, and the UK Medical Research Council (award 1324778), which was a PhD studentship administered by the University of Dundee. We thank Sarah Coulthurst, Sarah Murdoch, Matthias Trost, Rosie Clarke and everyone else on the chitinase secretion project at the University of Dundee (Jaeger Hamilton, Lucia Licandro Lado) for discussions, advice and reagents.

