## Supplementary Information for "Controlling and co-ordinating chitinase secretion in a *Serratia marcescens* population"

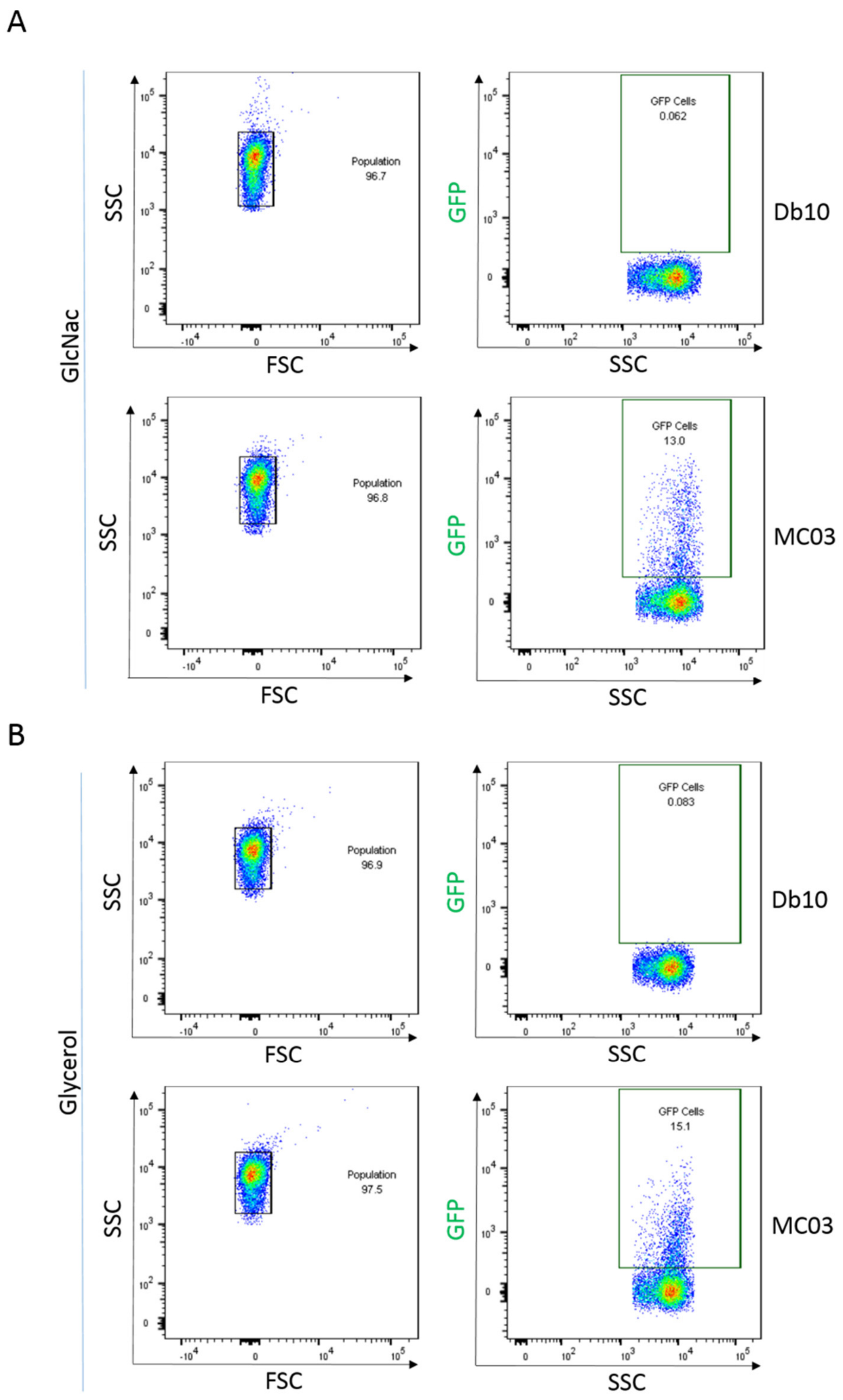


**Supp. Figure S1: A low proportion of cells express ϕ*gfp-chiC*.**

Samples were analysed on a LSR Fortessa (Becton Dickinson) flow cytometer and cells identified on the basis of forward scatter (FSC) and side scatter (SSC) and the GFP fluorescence quantified (488nm excitation, 530+/-30nm emission. MC03 and Db10 control population were compared. The non-GFP expressing Db10 control was used to evaluate background fluorescence and GFP positive cells identified on this basis. **A.** Cell growth in MM-GlcNAc displays 13% of GFP positive cells. **B.** Growth in MM-glycerol shows 15.1% of the population 'ON' for GFP fluorescence. Images were analysed and figures made using FlowJo v.10.1r7 version software.

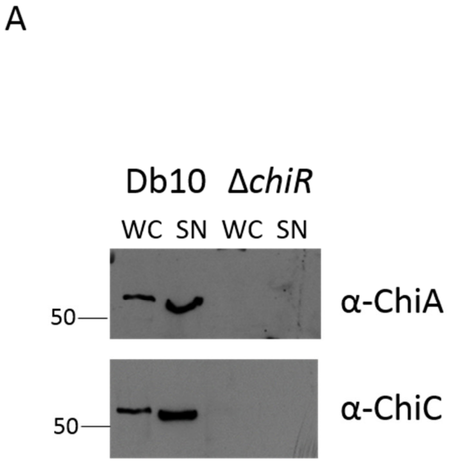


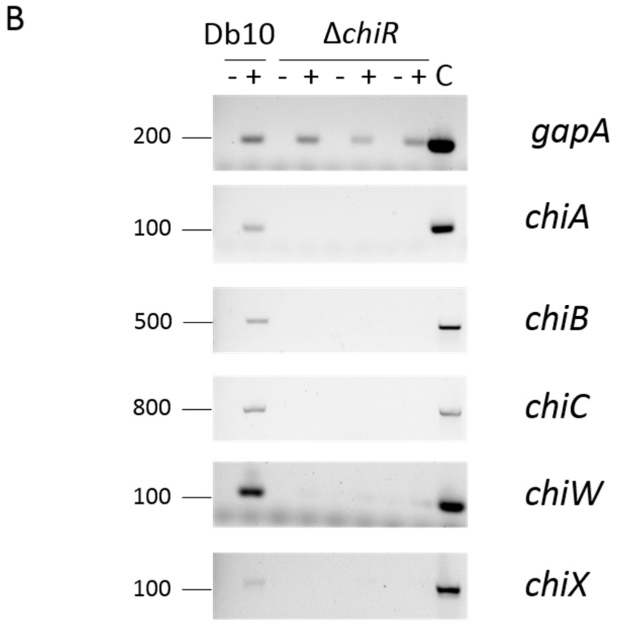


**Supp. Figure S2: ChiR is essential for chitinase transcription.**
**A.** Western blot of BC02 (Db10, Δ*chiR*). Cells were grown for 16 hours in rich medium prior to separation into 'WC' whole cells and 'SN' supernatant fractions **B.** Reverse transcriptase PCR of a Δ*chiR* strain**.** RNA was extracted from cells after 14 hours of growth in rich media and samples with Reverse transcriptase (+) and controls with no enzyme (-) were processed in four biological samples. *S. marcescens* Db10 genomic DNA was used as a PCR positive control (C). Primers were designed to amplify a 100-200 bp products of internal gene sequences. *gapA* = glyceraldehyde-3-phosphate dehydrogenase, thought to be constitutively expressed in *S. marcescens*.

**SUPP. Table S1: Total peptide and protein counts for proteomic samples.**

| | Biological Replicate | Peptides | Proteins |
|---|---|---|---|
| Δ*chiR* pBAD18-*chiR-TF* | 1 | 25856 | 2721 |
| | 2 | 24664 | 2706 |
| | 3 | 26163 | 2727 |
| | 4 | 25684 | 2721 |
| Δ*chiR* pBAD18 | 1 | 22585 | 2668 |
| | 2 | 24696 | 2722 |
| | 3 | 24498 | 2727 |
| | 4 | 25018 | 2731 |
| DB10 pBAD18 | 1 | 26202 | 2719 |
| | 2 | 25689 | 2720 |
| | 3 | 25823 | 2721 |
| | 4 | 25650 | 2719 |

Four biological replicates from each sample were analysed: the parental strain *S. marcescens* DB10 containing the pBAD18 empty vector, and the mutant train BC02 (Δ*chiR*) containing either pBAD18 or pBAD-ChiR3F.

**SUPP. TABLE S2**

| Protein ID | Protein Description | Score | -log10 (p-value) | log2 DB10 / ΔchiR |
|---|---|---|---|---|
| SMDB11_4243 | chiA endo-chitinase | 323.310 | 4.456 | 8.482 |
| SMDB11_2877 | cbp chitin-binding protein | 323.310 | 6.315 | 7.213 |
| SMDB11_0468 | chiC chitinase | 323.310 | 3.883 | 7.201 |
| SMDB11_2140 | narJ respiratory nitrate reductase 1 delta chain | 62.172 | 4.203 | 5.122 |
| SMDB11_4595 | nrdD anaerobic ribonucleoside-triphosphate reductase | 35.798 | 2.326 | 5.021 |
| SMDB11_2141 | narH respiratory nitrate reductase 1 beta chain | 101.670 | 2.303 | 4.871 |
| SMDB11_1290 | pepT peptidase T | 106.090 | 2.362 | 4.280 |
| SMDB11_1898 | putative formate acetyltransferase 3 | 14.341 | 5.394 | 4.195 |
| SMDB11_2706 | hypothetical protein | 12.717 | 1.820 | 4.056 |
| SMDB11_2156 | narK nitrate/nitrite transporter | 12.425 | 2.113 | 4.043 |
| SMDB11_2142 | narG respiratory nitrate reductase 1 alpha chain | 284.300 | 2.317 | 3.982 |
| SMDB11_3421 | aroP aromatic amino acid transport protein | 3.161 | 1.635 | 3.885 |
| SMDB11_2139 | narI respiratory nitrate reductase 1 gamma chain | 5.770 | 4.106 | 3.438 |
| SMDB11_0367 | ABC transporter, substrate-binding protein | 19.427 | 1.721 | 3.047 |
| SMDB11_3514 | gldA glycerol dehydrogenase, NAD | 10.945 | 2.811 | 2.782 |
| SMDB11_1746 | hypothetical protein | 6.996 | 1.461 | 2.766 |
| SMDB11_0226 | putative membrane protein | 15.811 | 3.296 | 2.694 |
| SMDB11_2216 | flgE flagellar hook protein | 8.325 | 2.089 | 2.605 |
| SMDB11_1941 | ompW outer membrane protein W | 76.975 | 2.047 | 2.474 |
| SMDB11_1547 | putative dethiobiotin synthetase | 20.915 | 1.954 | 2.464 |
| SMDB11_3842 | nirD nitrite reductase (NAD(P)H) small subunit | 7.452 | 1.374 | 2.380 |
| SMDB11_1691 | hypothetical protein | 3.789 | 2.363 | 2.252 |
| SMDB11_3070 | grcA autonomous glycyl radical cofactor | 279.000 | 2.177 | 2.198 |
| SMDB11_1202 | putative phosphoesterase | 5.198 | 2.270 | 2.072 |
| SMDB11_4639 | putative glycosyl transferase | 3.846 | 4.325 | 2.035 |
| SMDB11_4580 | hypothetical protein | 2.276 | 1.534 | 2.006 |
| SMDB11_1031A | rmf ribosome modulation factor | 6.883 | 1.332 | 1.914 |
| SMDB11_3841 | nirB nitrite reductase, large subunit, NAD(P)H-binding | 66.545 | 1.735 | 1.906 |
| SMDB11_2847 | two-component system sensor kinase | 4.363 | 2.353 | 1.831 |
| SMDB11_0368 | 50S ribosomal protein L31 type B | 7.526 | 1.882 | 1.793 |

| | | | | |
|---|---|---|---|---|
| SMDB11_0033 | putative exported protein | 19.625 | 1.725 | 1.786 |
| SMDB11_3870 | greB transcription elongation factor | 1.819 | 2.149 | 1.781 |
| SMDB11_2875 | chiB chitinase | 323.310 | 1.328 | 1.761 |
| SMDB11_0268 | putative methyltransferase | 12.510 | 1.324 | 1.719 |
| SMDB11_0954 | RpiR-family transcriptional regulator | 39.691 | 2.431 | 1.693 |
| SMDB11_0660 | TetR-family transcriptional regulator | 2.317 | 2.822 | 1.666 |
| SMDB11_4516 | methyl-accepting chemotaxis protein | 6.786 | 2.807 | 1.628 |
| SMDB11_2012 | putative metal-binding protein | 18.859 | 1.770 | 1.594 |
| SMDB11_0051 | sgrR DNA-binding transcriptional regulator | 4.686 | 1.998 | 1.592 |
| SMDB11_1249 | putative fimbrial chaperone protein | 18.886 | 2.730 | 1.587 |
| SMDB11_2639 | mhbM 3-hydroxybenzoate-6-hydroxylase | 2.347 | 1.413 | 1.485 |
| SMDB11_4603 | adenylation domain-containing protein | 9.345 | 1.645 | 1.465 |
| SMDB11_2043 | znuC zinc transporter subunit: ATP-binding component of ABC superfamily | 3.153 | 1.824 | 1.461 |
| SMDB11_0054 | putative transporter | 5.443 | 3.003 | 1.387 |
| SMDB11_2484 | putative exported protein | 3.722 | 2.348 | 1.365 |
| SMDB11_1248 | putative outer membrane fimbrial usher protein | 5.836 | 1.395 | 1.355 |
| SMDB11_0846 | dcuC anaerobic C4-dicarboxylate transporter | 13.229 | 2.582 | 1.275 |
| SMDB11_4682 | aldehyde dehydrogenase | 1.578 | 2.409 | 1.261 |
| SMDB11_4466 | frdB fumarate reductase iron-sulfur subunit | 67.299 | 1.774 | 1.245 |
| SMDB11_3472 | abgB aminobenzoyl-glutamate utilization protein B | 8.106 | 2.071 | 1.235 |
| SMDB11_2427 | putative polysaccharide deacetylase | 2.471 | 1.917 | 1.186 |
| SMDB11_4465 | frdC fumarate reductase subunit C | 7.071 | 2.313 | 1.158 |
| SMDB11_3219 | putative membrane protein | 6.562 | 1.315 | 1.142 |
| SMDB11_2229 | cheB chemotaxis reg: glutamatemethylesterase in two-component system with CheA | 21.107 | 2.146 | 1.086 |
| SMDB11_2500 | ABC transporter, substrate-binding protein | 7.686 | 2.365 | 1.050 |
| SMDB11_4453 | dcuA anaerobic C4-dicarboxylate transporter | 1.618 | 1.585 | 1.036 |
| SMDB11_0161 | csrA pleiotropic regulatory protein for carbonsource metabolism | 12.671 | 1.371 | 1.036 |
| SMDB11_3010 | pepB aminopeptidase B | 259.610 | 1.840 | 1.031 |
| SMDB11_4467 | frdA fumarate reductase, flavoprotein subunit | 205.280 | 2.224 | 1.018 |
| SMDB11_1124 | SMDB11_1124 hypothetical protein | 10.318 | 1.740 | -1.020 |
| SMDB11_3019 | iscR DNA-binding transcriptional repressor | 93.745 | 2.050 | -1.024 |

| | | | | |
|---|---|---|---|---|
| SMDB11_3358 | csaA putative protein secretion chaperone | 56.574 | 2.215 | -1.033 |
| SMDB11_2490 | putative alpha/beta hydrolase | 3.086 | 1.724 | -1.134 |
| SMDB11_3480 | exbB membrane spanning protein in TonB-ExbB-ExbD complex | 10.673 | 2.376 | -1.213 |
| SMDB11_3479 | exbD membrane spanning protein in TonB-ExbB-ExbD complex | 10.932 | 1.736 | -1.254 |
| SMDB11_3088 | putative membrane protein | 42.422 | 1.967 | -1.266 |
| SMDB11_1779 | fbpA iron(III) ABC transporter, substrate-binding protein | 140.310 | 1.991 | -1.271 |
| SMDB11_4406 | putative non-ribosomal peptide synthetase | 3.286 | 1.370 | -1.272 |
| SMDB11_4408 | fepA iron-enterobactin outer membrane transporter | 45.113 | 1.833 | -1.281 |
| SMDB11_4415 | entB isochorismatase | 28.976 | 2.479 | -1.341 |
| SMDB11_1411 | putative substrate-binding transport protein | 26.487 | 2.350 | -1.445 |
| SMDB11_3370 | putative pyridine nucleotide-disulphide oxidoreductase | 35.246 | 1.958 | -1.471 |
| SMDB11_1010 | putative exported protein | 2.848 | 2.885 | -1.728 |
| SMDB11_3364 | aldehyde dehydrogenase | 80.992 | 1.833 | -1.734 |
| SMDB11_3363 | putative thiamine pyrophosphate enzyme | 84.671 | 1.614 | -1.772 |
| SMDB11_4273 | wzxE O-antigen repeating unit flippase | 1.983 | 1.353 | -1.773 |
| SMDB11_3366 | putative short chain dehydrogenase | 59.975 | 1.307 | -1.776 |
| SMDB11_3969 | putative membrane protein | 10.515 | 1.736 | -1.790 |
| SMDB11_1061 | P13717 Nuclease | 79.794 | 2.141 | -1.889 |
| SMDB11_3376 | putative [2Fe-2S] protein | 4.652 | 2.094 | -1.905 |
| SMDB11_3368 | putative dioxygenase | 139.380 | 1.396 | -1.913 |
| SMDB11_3372 | IclR-family transcriptional regulator | 41.502 | 1.876 | -1.955 |
| SMDB11_3621 | hypothetical protein | 181.450 | 1.658 | -1.990 |
| SMDB11_1861 | putative membrane protein | 14.192 | 2.956 | -2.030 |
| SMDB11_3365 | putative dehydrogenase | 4.629 | 1.466 | -2.067 |
| SMDB11_3373 | putative short chain dehydrogenase | 63.264 | 1.621 | -2.071 |
| SMDB11_1933 | trpE component I of anthranilate synthase | 10.382 | 1.600 | -2.171 |
| SMDB11_3101 | putative membrane protein | 1.789 | 2.937 | -2.214 |
| SMDB11_0601 | putative bacteriocin | 11.449 | 1.373 | -2.313 |
| SMDB11_0658 | predicted protein | 9.207 | 1.468 | -2.567 |
| SMDB11_3374 | putative [2Fe-2S] protein | 61.956 | 1.406 | -2.712 |

| | | | | |
|---|---|---|---|---|
| SMDB11_3622 | hypothetical protein | 27.960 | 1.871 | -2.723 |
| SMDB11_0332 | major facilitator superfamily protein | 4.033 | 1.415 | -2.858 |
| SMDB11_2360 | tyrosine aminotransferase, tyrosine-repressible, PLP-dependent | 207.240 | 1.414 | -2.865 |
| SMDB11_2114 | phoE outer membrane pore protein | 6.529 | 1.324 | -3.106 |
| SMDB11_4409 | ferric-siderophore esterase | 8.256 | 2.889 | -3.543 |
| SMDB11_4411 | entF enterobactin synthase subunit F | 11.435 | 2.323 | -3.594 |
| SMDB11_1111 | hypothetical protein | 2.640 | 4.555 | -3.644 |

**SUPP. TABLE S3**

| Protein ID | Protein Description | Score | -log10 (p-value) | log2 ΔchiR +chiR/ΔchiR |
|---|---|---|---|---|
| SMDB11_4243 | chiA endo-chitinase | 323.310 | 4.616 | 9.942 |
| SMDB11_2876 | chiR LysR-family transcriptional regulator | 88.710 | 4.946 | 8.218 |
| SMDB11_2877 | cbp chitin-binding protein | 323.310 | 5.792 | 7.732 |
| SMDB11_0468 | chiC chitinase | 323.310 | 3.477 | 6.310 |
| SMDB11_2875 | chiB chitinase | 323.310 | 4.535 | 5.281 |
| SMDB11_4595 | nrdD anaerobic ribonucleoside-triphosphate reductase | 35.798 | 2.013 | 4.414 |
| SMDB11_2873 | putative phage-related lysin (ChiX) | 5.093 | 3.538 | 4.344 |
| SMDB11_2140 | narJ respiratory nitrate reductase 1 delta chain | 62.172 | 2.587 | 4.206 |
| SMDB11_2872 | putative phage-related exported protein (ChiY) | 19.091 | 4.499 | 3.995 |
| SMDB11_2141 | narH respiratory nitrate reductase 1 beta chain | 101.670 | 1.631 | 3.674 |
| SMDB11_3421 | aroP aromatic amino acid transport protein | 3.161 | 1.339 | 3.252 |
| SMDB11_1290 | pepT peptidase T | 106.090 | 1.569 | 3.184 |
| SMDB11_2156 | narK nitrate/nitrite transporter | 12.425 | 1.718 | 3.116 |
| SMDB11_0041 | rluA 23S rRNA/tRNA pseudouridine synthase A | 1.952 | 2.254 | 3.038 |
| SMDB11_1746 | hypothetical protein | 6.996 | 2.440 | 3.013 |
| SMDB11_2142 | narG respiratory nitrate reductase 1 alpha chain | 284.300 | 1.724 | 2.991 |
| SMDB11_2139 | narI respiratory nitrate reductase 1 gamma chain | 5.770 | 3.584 | 2.699 |
| SMDB11_3514 | gldA glycerol dehydrogenase, NAD | 10.945 | 2.667 | 2.415 |
| SMDB11_2910 | putative transcriptional regulator | 6.763 | 2.514 | 2.069 |
| SMDB11_4639 | putative glycosyl transferase | 3.846 | 3.693 | 2.049 |
| SMDB11_2216 | flgE flagellar hook protein | 8.325 | 1.746 | 1.975 |
| SMDB11_1202 | putative phosphoesterase | 5.198 | 2.074 | 1.964 |
| SMDB11_0268 | putative methyltransferase | 12.510 | 1.539 | 1.838 |
| SMDB11_1547 | putative dethiobiotin synthetase | 20.915 | 1.527 | 1.832 |
| SMDB11_2847 | two-component system sensor kinase | 4.363 | 1.943 | 1.705 |
| SMDB11_1941 | ompW outer membrane protein W | 76.975 | 1.326 | 1.690 |
| SMDB11_0660 | TetR-family transcriptional regulator | 2.317 | 2.389 | 1.448 |
| SMDB11_0723 | putative acetyltransferase | 4.806 | 1.577 | 1.439 |

| | | | | |
|---|---|---|---|---|
| SMDB11_4105 | mobB molybdopterin-guanine dinucleotide biosynthesis protein B | 2.954 | 1.460 | 1.420 |
| SMDB11_3674 | fis DNA-binding protein Fis | 2.084 | 1.394 | 1.252 |
| SMDB11_3731 | wzi putative surface assembly of capsule protein | 4.800 | 1.319 | 1.190 |
| SMDB11_0780 | putative exported protein | 8.326 | 1.307 | 1.163 |
| SMDB11_3456 | type VI secretion system, effector protein Hcp-like | 6.996 | 1.888 | 1.092 |
| SMDB11_2744 | tctA tricarboxylic transport protein | 2.455 | 1.432 | 1.025 |
| SMDB11_4611 | putative oxidoreductase | 37.985 | 2.391 | -1.004 |
| SMDB11_2450 | isochorismatase | 11.940 | 2.811 | -1.011 |
| SMDB11_2176 | efeO ferrous iron transport periplasmic protein | 103.440 | 2.951 | -1.021 |
| SMDB11_4702 | hpcE2 4-hydroxyphenylacetate degradation protein | 56.176 | 3.099 | -1.022 |
| SMDB11_1267 | hemK protein methyltransferase | 3.102 | 1.337 | -1.025 |
| SMDB11_0829 | mglB methyl-galactoside transporter subunit | 270.920 | 3.932 | -1.055 |
| SMDB11_1323 | hypothetical protein | 199.350 | 2.620 | -1.082 |
| SMDB11_2932 | putative phosphate ABC transporter, permease protein | 3.448 | 1.533 | -1.091 |
| SMDB11_1779 | fbpA iron(III) ABC transporter, substrate-binding protein | 140.310 | 2.351 | -1.205 |
| SMDB11_4710 | hpaB 4-hydroxyphenylacetate 3-monooxygenaseoxygenase component | 123.580 | 2.800 | -1.206 |
| SMDB11_1411 | SMDB11_1411 putative substrate-binding transport protein | 26.487 | 2.565 | -1.245 |
| SMDB11_4706 | hpcG 2-oxo-hepta-3-ene-1,7-dioic acid hydratase | 35.764 | 2.782 | -1.266 |
| SMDB11_0659 | predicted protein | 3.399 | 2.038 | -1.305 |
| SMDB11_3479 | exbD membrane spanning protein in TonB-ExbB-ExbD complex | 10.932 | 1.703 | -1.318 |
| SMDB11_4408 | fepA iron-enterobactin outer membrane transporter | 45.113 | 2.257 | -1.320 |
| SMDB11_4415 | entB isochorismatase | 28.976 | 2.630 | -1.338 |
| SMDB11_2059 | phage protein | 43.727 | 1.825 | -1.453 |
| SMDB11_4414 | entE enterobactin synthase subunit E | 7.318 | 1.734 | -1.511 |
| SMDB11_2533 | putative ribose ABC transporter,substrate-binding protein | 7.722 | 2.843 | -1.529 |
| SMDB11_3370 | putative pyridine nucleotide-disulphide oxidoreductase | 35.246 | 2.027 | -1.620 |
| SMDB11_1010 | putative exported protein | 2.848 | 2.041 | -1.630 |
| SMDB11_3622 | hypothetical protein | 27.960 | 1.789 | -1.639 |
| SMDB11_3366 | putative short chain dehydrogenase | 59.975 | 2.080 | -1.639 |
| SMDB11_3094 | hypothetical protein | 75.723 | 2.988 | -1.653 |
| SMDB11_1233 | ABC transporter, ATP-binding protein | 1.667 | 2.088 | -1.690 |

| | | | | |
|---|---|---|---|---|
| SMDB11_2534 | putative zinc-binding alcohol dehydrogenase | 19.894 | 2.893 | -1.714 |
| SMDB11_4712 | ampH penicillin-binding protein | 4.808 | 1.519 | -1.778 |
| SMDB11_3372 | IclR-family transcriptional regulator | 41.502 | 2.455 | -1.790 |
| SMDB11_0347 | putative exported protein | 2.620 | 1.675 | -1.793 |
| SMDB11_1061 | P13717 Nuclease | 79.794 | 2.308 | -1.849 |
| SMDB11_3363 | putative thiamine pyrophosphate enzyme | 84.671 | 2.677 | -1.885 |
| SMDB11_3373 | putative short chain dehydrogenase | 63.264 | 2.568 | -1.892 |
| SMDB11_3376 | putative [2Fe-2S] protein | 4.652 | 3.009 | -1.914 |
| SMDB11_3364 | aldehyde dehydrogenase | 80.992 | 2.651 | -1.959 |
| SMDB11_3368 | putative dioxygenase | 139.380 | 2.335 | -2.015 |
| SMDB11_3375 | hypothetical protein | 8.688 | 2.052 | -2.044 |
| SMDB11_2360 | tyrosine aminotransferase, tyrosine-repressible, PLP-dependent | 207.240 | 2.313 | -2.073 |
| SMDB11_1721 | TonB-dependent copper receptor | 32.484 | 1.797 | -2.077 |
| SMDB11_2915 | transcriptional regulator | 2.707 | 1.481 | -2.107 |
| SMDB11_3371 | dioxygenase superfamily protein | 222.370 | 1.918 | -2.271 |
| SMDB11_0675 | enhancin | 59.408 | 2.221 | -2.291 |
| SMDB11_4160 | putative lipoprotein | 3.933 | 1.840 | -2.303 |
| SMDB11_4409 | ferric-siderophore esterase | 8.256 | 1.329 | -2.316 |
| SMDB11_4407 | putative iron transport protein | 20.051 | 1.478 | -2.438 |
| SMDB11_0658 | predicted protein | 9.207 | 2.434 | -2.512 |
| SMDB11_3374 | putative [2Fe-2S] protein | 61.956 | 2.381 | -2.518 |
| SMDB11_3365 | putative dehydrogenase | 4.629 | 1.748 | -2.550 |
| SMDB11_1350 | putative chorismate mutase precursor | 2.020 | 3.795 | -3.000 |
| SMDB11_1111 | hypothetical protein | 2.640 | 3.707 | -3.089 |
| SMDB11_1738 | ankyrin repeat-containing protein | 26.271 | 1.792 | -3.296 |
| SMDB11_0332 | major facilitator superfamily protein | 4.033 | 2.051 | -3.326 |
| SMDB11_3367 | putative alpha/beta hydrolase | 12.424 | 5.978 | -4.541 |
